# Ultrafast and Accurate Selection of High-Quality Protein Complex Models from Large-Scale Prediction

**DOI:** 10.64898/2026.09.26.754618

**Authors:** Fang Liang, Lei Xie, Enjia Ye, Zhengri Pan, Tiantian Xu, Haodong Wang, Pengcheng Wei, Jun Liu, Guijun Zhang

**Affiliations:** College of Information Engineering, Zhejiang University of Technology, Hangzhou 310023, China; Zhejiang Key Laboratory of Intelligent Perception and Control for Complex Systems, Zhejiang University of Technology, 288 Liuhe Road, Hangzhou 310023, China; Cancer Science Institute of Singapore, National University of Singapore, Singapore, Singapore

**Keywords:** Large-scale protein structural model selection, Estimated model accuracy, Predictor-derived confidence, Ultrafast shape recognition, Structural consensus

## Abstract

In the post-AlphaFold era, advances in protein complex structure prediction have enabled the generation of thousands of candidate models per target, making the accurate and efficient selection of high-quality models from large candidate pools a critical challenge. Existing model-selection approaches commonly employ estimated model accuracy (EMA) methods, which include single-model and consensus-based methods. The former lack cross-model comparative information, whereas the latter often incur high computational costs from pairwise structural alignments and are influenced by candidate-pool quality. Here, we present DeepUMQA-Selection, a two-stage framework combining single-model accuracy estimators for preliminary selection with Ultrafast Shape Recognition (USR)-based structural alignment for consensus selection. Preliminary selection enriches the candidate pool with high-quality models and reduces subsequent structural comparisons, while USR-based structural alignment enables efficient consensus selection without explicit structural superposition. On the retrospective CASP16 QMODE3 benchmark dataset, DeepUMQA-Selection outperformed all participating methods and predictor-derived confidence metrics, as measured by mean Top-5 weighted penalty. The advantage was particularly pronounced for heteromeric targets, with a 13.9% improvement in model-selection performance relative to the best-performing CASP16 participating method. Furthermore, a confidence-assisted strategy enabled the framework to accommodate larger protein complexes while maintaining superior model-selection performance over competing methods, supporting its scalability. Runtime benchmarking showed that the proposed USR-based structural alignment achieved a substantial 317-fold speedup over US-align for candidate pools containing 1,000 models, demonstrating its computational efficiency for large-scale model selection. Our results demonstrate that DeepUMQA-Selection enables accurate and computationally efficient selection of high-quality protein complex models from large-scale candidate pools.

## 1 Introduction

Recent advances in deep learning-based protein structure prediction, exemplified by the AlphaFold series[1-3] and emerging approaches such as Protenix-v1[4], Boltz-2[5], and Chai-1[6], now make it feasible to generate thousands of candidate structures for a single target. Such advances have fundamentally shifted the field’s core bottleneck from model generation to model selection. Most existing approaches, however, rely on confidence metrics derived from the modeling process itself. These metrics are not only tightly coupled with specific architectural designs but also exhibit substantial calibration biases on challenging targets[7,8], rendering them unreliable for robust, cross-method selection at scale. This pressing gap underscores the urgent need for a model-agnostic, third-party framework capable of accurately and efficiently prioritizing the best models from large-scale candidate pools, thereby providing a critical step toward realizing the full potential of structural prediction in drug discovery and beyond.

Model-agnostic estimated model accuracy (EMA) methods have long served as the primary tools for conventional model quality estimation and ranking[9]. However, these methods were largely designed for small-scale model sets and fall into three categories based on how they leverage structural information. Single-model approaches (e.g., DeepAccNet[10], VoroIF-GNN[11], DeepUMQA[12-14]) score each model independently, offering high computational efficiency but often focusing on specific quality metrics and lacking the comparative signals from other candidates that are critical for reliable ranking[7,9]. Quasi-single-model methods (e.g., MIEnsemble-server[8], ModFOLDdock2S[15]) partially address this by incorporating limited information from internally generated models, striking a balance between efficiency and consensus. Consensus-based methods (e.g., ModFOLD[16-18], MULTICOM[19-21], VoroIF-jury[22], DeepUMQA-X[23]) derive quality estimates from structural agreement across candidates and generally achieve superior ranking performance at the cost of exhaustive pairwise alignments that become prohibitively expensive as model pools grow[9]. Most EMA methods, though model-agnostic, were designed for small-scale model sets, with single-model approaches lacking the comparative context needed to rank reliably among thousands of candidates and consensus-based methods demanding exhaustive alignments that become computationally prohibitive at scale[24].

Recognizing the emerging challenge of large-scale model selection, the 16th Critical Assessment of protein Structure Prediction (CASP16), a community-wide blind assessment, introduced the QMODE3 category in 2024 to evaluate whether existing methods can reliably identify the best models from massive candidate pools[7]. Participants were tasked with selecting the Top-5 models predicted to be of the highest quality from pools generated by MassiveFold[25,26], a scalable AlphaFold-based framework that enables massively parallelized inference to generate thousands of models per target. The selected models were assessed using a covariance-weighted penalty score[7], an official CASP16 metric where lower values indicate better selections. This score integrates multiple quality metrics, such as TM-score[27], DockQ[28], and lDDT[29], and accounts for their inherent correlations, thereby avoiding redundant contributions and yielding a more accurate reflection of selection quality. Notably, most participating strategies were directly migrated from existing EMA approaches. The results showed that these EMA methods could not match the performance of predictor-derived confidence metrics (ipTM and ipTM+pTM) on heteromeric targets. On average, the best-selected model had a true rank of 147 by TM-score[27] or 62 by DockQ-ave[9,28], possibly confirming previous observations that current EMA methods still struggle to identify the top-quality models available in the prediction model pools[7].

In summary, the CASP16 blind assessment results reveal a fundamental dilemma in large-scale model selection, where accuracy and efficiency are inherently difficult to reconcile. Single-model EMA methods are computationally efficient but lack the comparative context required for reliable ranking. Consensus-based EMA methods offer the comparative signals essential for reliable ranking, yet their reliance on pairwise structural alignments becomes computationally prohibitive at the scale of thousands of models. The composition of the model pool introduces further complications, since when high-quality models are scarce or structural biases pervade the candidate set, consensus signals can become misleading, substantially increasing the risk of selecting suboptimal models[30,31]. These observations give rise to two interconnected challenges: how to reduce the number of models requiring pairwise comparison through effective prefiltering, and how to perform each comparison efficiently without resorting to computationally expensive structural superposition (e.g., US-align[32]).

To address these challenges, we introduce DeepUMQA-Selection, a two-stage framework designed for reliable and efficient prioritization of protein complex models from large-scale prediction pools. The first stage employs two complementary single-model quality estimators to perform an initial screening of the candidate pool. This step not only reduces the number of models requiring subsequent comparison but also mitigates potential biases arising from overrepresented conformational clusters, thereby enriching the pool with high-quality candidates. The second stage applies Ultrafast Shape Recognition (USR)[12-14,33]-based consensus alignment to efficiently rank the retained models, bypassing the need for computationally expensive structural superposition while preserving the comparative signals essential for reliable selection. Together, this two-stage design enables DeepUMQA-Selection to achieve both high accuracy and computational efficiency, with the former validated on the CASP16 QMODE3 benchmark[7] and the latter assessed against the widely used structural alignment tool US-align[32].

## 2. Materials and methods

### 2.1 Dataset

To evaluate DeepUMQA-Selection for large-scale protein complex model selection, we used the MassiveFold-generated candidate model pools from the CASP16 QMODE3 assessment as the benchmark. QMODE3 was designed to evaluate the ability of model quality assessment methods to select Top-5 models predicted to be of the highest quality from large pools of protein complex models. The benchmark comprised 36 protein complex targets, including 15 homomerics and 21 heteromerics, of which 8 were antibody–antigen complexes. In total, 259,835 candidate models were included in the evaluation (**Supplementary Table S1**). Among these targets, 23 complexes had sequence lengths of ≤1,500 residues, including 14 heteromeric and 9 homomeric complexes (**Supplementary Table S2**). The remaining 13 complexes, each containing >1,500 residues, comprised 7 heteromeric and 6 homomeric complexes (**Supplementary Table S3**).

### 2.2 Overview of DeepUMQA-Selection

DeepUMQA-Selection is a two-stage framework for selecting high-quality models from large-scale protein complex prediction pools (**Figure 1**). In the first stage, the candidate pool undergoes an initial screening to reduce its size, thereby enriching it with high-quality candidates. In the second stage, Ultrafast Shape Recognition (USR)-based consensus alignment is applied to efficiently re-rank the retained models, bypassing computationally expensive structural superposition while preserving the comparative signals essential for reliable selection, with the Top-*K* models selected as the final output.

**Figure 1.**
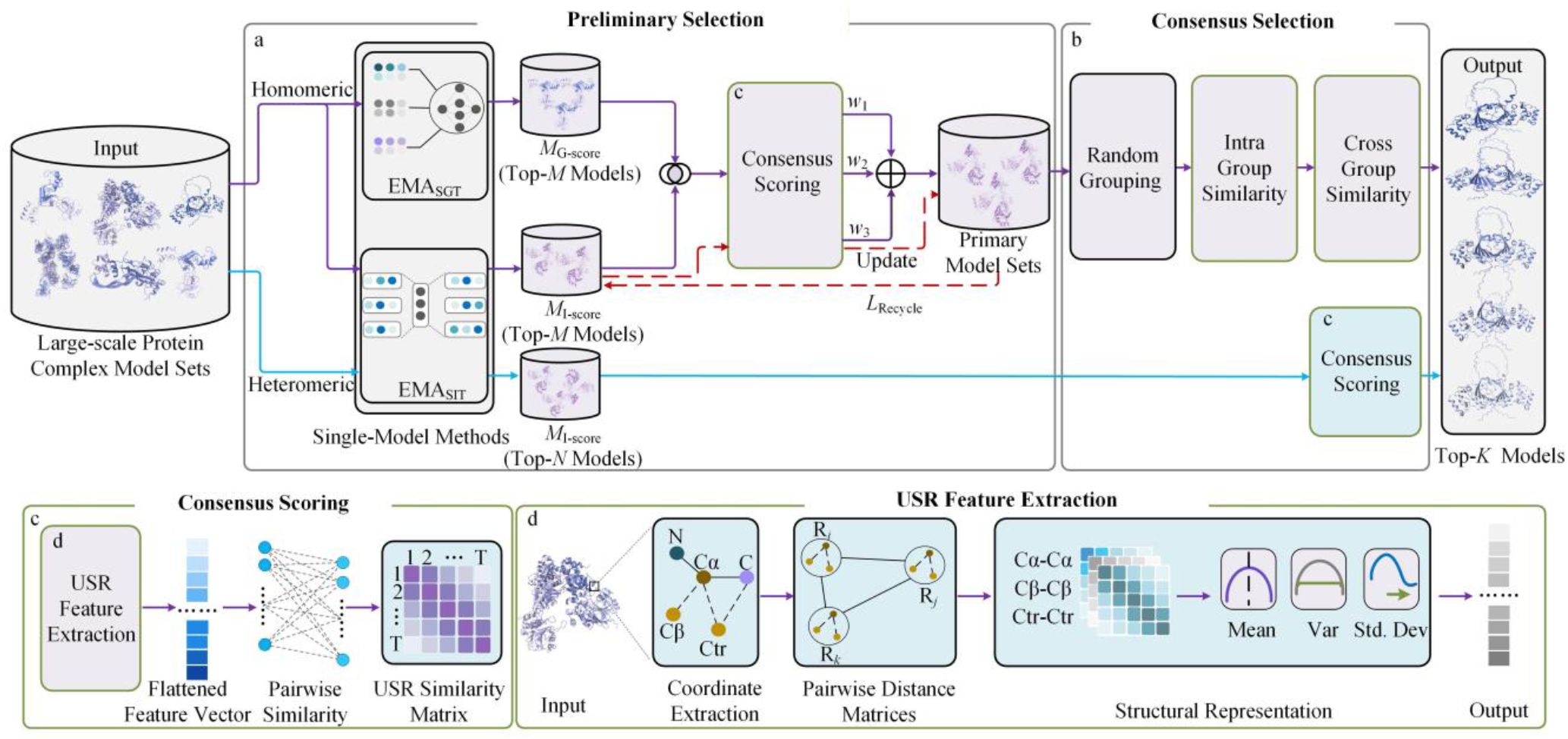
Overview of DeepUMQA-Selection. For each protein complex target, DeepUMQA-Selection processes a large candidate model pool through two stages: preliminary selection and consensus selection to identify the final Top-*K* high-quality models. **(a) Preliminary selection**. For homomeric complexes, all candidate models are independently evaluated by two complementary single-model EMA methods, EMA_SGT_ (MViewEMA) and EMA_SIT_ (DeepUMQA-Global). The Top-*M* models from each ranking are retained, and their intersection is subjected to USR-based consensus scoring. The normalized EMA_SGT_, EMA_SIT_, and USR consensus scores are combined using weights *w*_1_, *w*_2_, and *w*_3_, respectively, to construct the primary model set, which is subsequently refined through iterative replacement for up to *L*_Recycle_ rounds. For heteromeric complexes, EMA_SIT_ is used for preliminary selection: the Top-*N* models form the reference set, while models with EMA_SIT_ scores above the target-specific median form the selection pool for subsequent consensus ranking. **(b) Consensus selection**. For homomeric complexes, the refined primary model set is randomly partitioned into groups and ranked using hierarchical intra-group and cross-group USR consensus. For heteromeric complexes, each model in the selection pool is ranked according to its mean USR similarity to the reference models. The Top-*K* models are returned as the final selection. **(c) Consensus scoring**. Model-level USR descriptors are flattened into one-dimensional vectors, and the Euclidean distance between each pair of vectors is transformed into a USR similarity score used for consensus ranking. **(d) USR feature extraction**. Residue-level C_α_, C_β_, and centroid coordinates (Ctr) define three coordinate representations. For each representation, distance distributions associated with the residue itself, the residue farthest from it, and the residue farthest from that farthest residue are summarized by their mean (Mean), variance (Var), and standard deviation (Std.Dev), yielding a nine-dimensional descriptor. The three descriptors are concatenated to form a 27-dimensional residue-level USR descriptor. Detailed parameter settings are provided in **Supplementary Table S4.**

While the above two-stage strategy applies to all targets, the specific implementations differ between homomeric and heteromeric complexes, reflecting their distinct accuracy distributions, with homomeric pools typically exhibiting quality scores concentrated near the high end with limited variance, whereas heteromeric pools show greater variability and long tails extending into the low-quality range (**Supplementary Figure S1**). This design distinction ensures that the consensus comparison is tailored to the quality distribution of each target type, and coincides with the separate ranking schemes adopted in the CASP16 assessment[7].

Specifically, for homomeric complexes, where candidate models share highly similar global folds and high-quality models are relatively abundant, the pipeline employs both EMA_SGT_ (MViewEMA[34]) for global topology assessment and EMA_SIT_ (DeepUMQA-Global[35]) for interface topology evaluation in the first stage (see **Section 2.6** for details). Their complementary assessments are intersected to construct a primary model set, which is then refined through an iterative replacement procedure to further concentrate high-quality candidates. In the second stage, the refined set is randomly divided into eight groups and subjected to hierarchical USR-based consensus ranking, where models are first ranked within each group and the group-level rankings are then integrated to generate a global ranking, enabling fine-grained discrimination among structurally similar models, a critical capability given the limited conformational spread in homomeric pools.

For heteromeric complexes, where model pools exhibit greater conformational diversity and high-quality models are often scarce, the first stage uses only single-model EMA_SIT_ for interface topology evaluation to identify a reference set of top-scoring models, which serve as reliable anchors representing the most plausible predictions. Meanwhile, all remaining models above the median score are retained as the selection pool. In the second stage, rather than performing exhaustive pairwise comparisons across the entire pool, which would be computationally expensive and prone to signal dilution, each model in the selection pool is ranked by its average USR similarity to the reference set. This reference-anchored strategy ensures that consensus signals are driven by high-quality references, effectively preventing low-quality or structurally divergent models from distorting the ranking, a critical advantage when high-quality models represent only a small fraction of the pool.

### 2.3 Ultrafast Shape Recognition (USR)-Based Structural Alignment

Consensus-based model selection depends critically on the efficient comparison of structural similarity among candidate models. Classical structural alignment methods, such as US-align[32], achieve high accuracy through iterative superposition to determine optimal residue correspondences and derive alignment-based similarity scores; however, their exhaustive pairwise application becomes computationally prohibitive when model pools expand to thousands of candidates. To alleviate this scalability bottleneck, we leverage the residue-level USR descriptors originally developed in our DeepUMQA series[12-14], where they served as input features to neural networks for estimating per-residue model accuracy, such as lDDT[29]. Here, we repurpose these descriptors as geometric fingerprints for pairwise structural comparison in consensus-based selection. Specifically, each residue is characterized by distance distributions from three reference points: itself, the residue farthest from it, and the residue farthest from that farthest residue. These distributions are derived from three coordinate representations, namely C_α_ atoms, reconstructed C_β_ atoms, and residue centroids, to form a rotationally and translationally invariant fingerprint for each residue. Model-level similarity can then be computed directly as the Euclidean distance between these fingerprints, bypassing explicit structural superposition. This adaptation preserves the discriminatory capability needed for consensus ranking while reducing the computational cost by orders of magnitude relative to exhaustive US-align comparisons, making accurate and scalable model selection feasible for large-scale candidate pools. Detailed pseudocode for the USR-based structural alignment procedure is provided in **Supplementary Algorithm 1**.

#### 2.3.1 Residue-level USR descriptors

For each residue *i* in a protein structure model, three representative coordinate points were extracted, including the C_*α*_, reconstructed C_β_, and residue centroid (C_centroid_). The residue centroid was calculated as the geometric center of all heavy atoms within the residue.

For each coordinate type *p* (*p* ∈{C_α_,C_β_,C_centroid_}) of residue *i*, two reference residues were identified based on Euclidean distances. The first reference residue 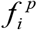 was defined as the residue whose corresponding coordinate point had the maximum Euclidean distance from that of residue *i*, and the second reference residue 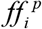 was identified as the residue with the largest Euclidean distance from 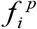.

Based on these three characteristic residues (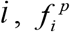, and 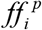), the Euclidean distance distributions from each characteristic residue to all residues were calculated (excluding itself). The resulting distance distributions were then characterized by three statistical descriptors, including the mean (*m*), variance (*v*), and standard deviation (*s*). Therefore, each coordinate type generated a 9-dimensional descriptor:

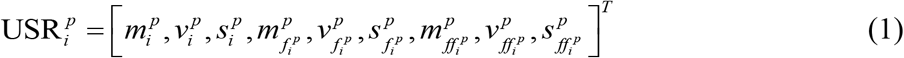

where *p* ∈{C_α_,C_β_,C_centroid_}. By combining the descriptors derived from the three representative coordinates, the final residue-level USR descriptor for residue *i* was obtained as a 27-dimensional vector:

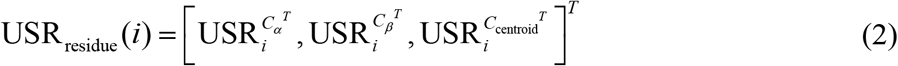

#### 2.3.2 Model-level USR descriptors

Based on the residue-level USR descriptors, a model-level USR descriptor was constructed to characterize the overall geometric features of each protein structure model. For a protein structure model containing *N* residues, the residue-level descriptors were sequentially arranged according to their amino acid sequence order:

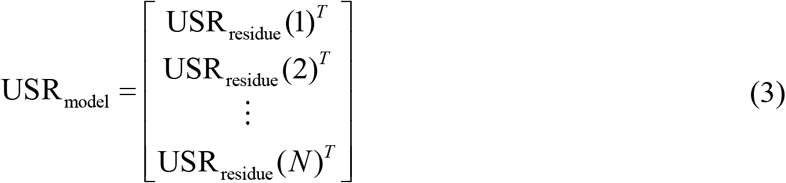

where USR_residue_ (*i*) represents the 27-dimensional USR descriptor of residue *i*. Therefore, the resulting model-level USR descriptor was represented as a *N* × 27 real-valued matrix, where each row corresponds to one residue and each row vector represents the 27-dimensional residue-level USR descriptor of the corresponding residue.

#### 2.3.3 USR similarity score

Based on the model-level USR descriptors, a pairwise USR similarity score was calculated to quantify the structural similarity between two protein models. Given two candidate models *A* and *B*, their corresponding model-level USR descriptors were first generated as 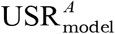 and 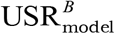, respectively.

To enable direct comparison between the two model-level descriptors, the *N* × 27 USR matrices were flattened into one-dimensional vectors while preserving the residue order. The flattened descriptors of the two models were denoted as: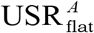 and 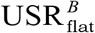, respectively, each represented as a real-valued vector of length 27*N*.

The Euclidean distance between the two flattened USR descriptors was then calculated to measure their geometric difference. Finally, the USR similarity score between models *A* and *B* was defined as:

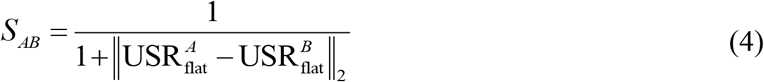

where *S*_*AB*_ represents the structural similarity between models *A* and *B*. *A* higher *S*_*AB*_ value indicates a smaller difference between their USR descriptors and therefore greater structural similarity.

### 2.4 Consensus selection for homomeric complexes

Homomeric candidate pools often contain many models with similar global architectures, making exhaustive structural-consensus analysis computationally demanding and reducing the discrimination provided by global similarity alone. We therefore developed a progressive consensus-selection pipeline that combines two independent single-model quality estimators with USR-based structural consensus. Single-model assessment first restricts consensus evaluation to quality-enriched candidates, while the use of two estimators reduces dependence on method-specific inductive biases and systematic ranking errors. The retained candidates are subsequently refined through iterative primary-set updating and hierarchical grouped consensus selection. The complete candidate pool is used as input, and the Top-*K* models in the final consensus ranking are returned. All parameter settings are provided in **Supplementary Table S4**.

#### 2.4.1 Consensus scoring using USR-Based Structural Alignment

All candidate models were independently evaluated by EMA_SGT_ and EMA_SIT_. The Top-*M* models from each ranking were retained, and their intersection was used for consensus scoring. This intersection restricts subsequent structural comparisons to candidates supported by both single-model estimators, thereby reducing the influence of estimator-specific ranking errors and the computational cost of pool-level analysis.

For each model in the intersection with a valid USR representation, the consensus score was calculated as its mean USR similarity to all other retained models, excluding self-comparison. This score measures the structural support received by a model within the quality-filtered candidate set and complements the model-specific evidence provided by the two estimators.

The EMA_SGT_, EMA_SIT_ and USR consensus scores were independently rescaled to [0,1] by min– max normalization and integrated as:

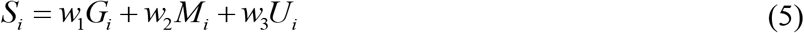

where *G*_*i*_, *M*_*i*_ and*U*_*i*_ denote the normalized EMA_SGT_, EMA_SIT_ and USR consensus scores of model *i*, respectively, and *w*_1_, *w*_2_ and *w*_3_ are the corresponding weights. Models were ranked in descending order of *S*_*i*_, and the highest-ranked candidates were retained as the initial primary model set.

#### 2.4.2 Primary model set mini-rollout update

The initial primary model set is constructed from a static combination of single-model quality estimates and USR consensus and may therefore retain models near the selection boundary that score acceptably on individual estimators but receive limited structural support from the consensus. To refine its composition, we introduced a mini-rollout update that iteratively replaces models with limited structural support and progressively concentrates the primary model set within a stable consensus region.

A broader quality-filtered reference set was first constructed from the original candidate set using a predefined single-model quality criterion. Let *R* denote the number of models replaced at each iteration and *L*_Recycle_ the maximum number of iterations. At each iteration, every model in the current primary model set was scored by its mean USR similarity to the reference set. The *R* models with the lowest scores were designated for removal. Models in the reference set that were not already included in the current primary model set were independently scored by their mean USR similarity to the current primary model set, and the *R* highest-scoring candidates were selected as replacements. The selected candidates replaced the designated models while the size of the primary model set remained unchanged. Detailed pseudocode for the consensus-driven dynamic pool iterative optimization procedure is provided in **Supplementary Algorithm 2**.

The updated primary model set was used as the input to the next iteration. The procedure terminated when *L*_Recycle_ iterations had been completed or when the composition of the primary model set remained unchanged between successive iterations. The resulting refined primary model set was subsequently used for hierarchical consensus selection.

#### 2.4.3 Hierarchical Consensus Selection

The refined primary set was subjected to hierarchical consensus selection to reduce repeated full-set comparisons and limit interference from lower-confidence candidates. Models were first randomly divided into *G* groups, allowing consensus calculations to be performed within smaller working sets before the resulting local rankings were integrated globally.

Within each group, models were independently ranked by EMA_SGT_ and EMA_SIT_. Models at corresponding rank positions in the two rankings were progressively appended to a cumulative set. Let *A*_*t*_ denote the models included after polling round *t*. The current consensus score of model *i* ∈ *A*_*t*_ was calculated as

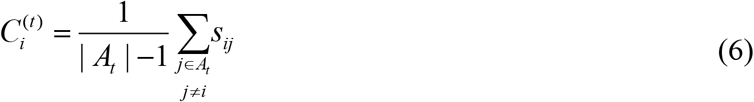

where |*A*_*t*_| denotes the number of models in *A*_*t*_, and *s*_*ij*_ is the USR similarity between model *i* and *j*. As lower-ranked models are progressively introduced, they may weaken a previously established consensus signal. The maximum score observed for each model across all relevant polling rounds was therefore retained:

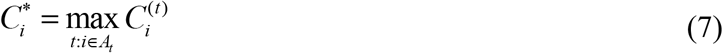

Models in each group were ranked according to 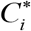 to generate intra-group consensus rankings. These rankings were then combined using the same cumulative procedure: models at corresponding positions from all group-level rankings were progressively appended to a global set, and their consensus scores were recalculated. The maximum cross-group consensus score was retained as the final score for each model. All models were ranked by their final consensus scores, and the Top-*K* models were returned as the final selection. The complete algorithm and pseudocode are provided in **Supplementary Algorithm 3**.

### 2.5 Consensus selection for heteromeric complexes

Compared with homomeric complexes, candidate models of heteromeric complexes generally exhibit greater structural diversity due to more complex inter-chain interactions and larger variations in predicted interface arrangements. This increased heterogeneity among candidate structures can make consensus-based selection more susceptible to interference from low-quality models when the entire model pool is considered. To address this challenge, we adopted a reference-guided consensus ranking strategy for heteromeric complexes.

First, we employed EMA_SIT_ to estimate the accuracy of all candidate models. The models were ranked in descending order according to their predicted scores, and the Top-*N* models were selected to construct a reference model set. Meanwhile, models with EMA_SIT_ scores higher than the median score of the corresponding candidate pool were retained as the selection pool, thereby reducing the influence of low-quality models on subsequent consensus-based ranking. For each model in the selection pool, its USR-based similarity scores with all reference models were calculated, and the average similarity score was used as the reference-guided consensus score. The models were then ranked according to these consensus scores, and the Top-*K* models were selected as the final selection.

This strategy integrates single-model preliminary selection with consensus-based ranking, enabling consensus evaluation to focus on a quality-enriched subset rather than the entire heterogeneous model pool. By restricting structural similarity calculations to the selection models and reference models, the proposed approach substantially reduces the computational burden in large-scale model selection while maintaining effective identification of high-quality models.

### 2.6 Single-model accuracy estimators

In the preliminary selection stage, we employed two complementary pretrained single-model quality estimators, EMA_SGT_ and EMA_SIT_, to provide model-agnostic assessments of individual protein complex models. Specifically, EMA_SGT_ and EMA_SIT_ correspond to our previously developed methods, MViewEMA[34] and DeepUMQA-Global[35], respectively. MViewEMA estimates the global accuracy of protein complex models by integrating complementary structural representations at the macro-, meso-, and micro-views[34]. These representations are derived solely from the input structure, without requiring templates, multiple sequence alignments, protein language models, or information from other candidate models. DeepUMQA-Global evaluates protein complex models using a structure– sequence cross-consistency mechanism that quantifies the bidirectional compatibility between the input sequence and the predicted three-dimensional structure, while also incorporating interface-aware structural information[35]. DeepUMQA-Global provides global quality assessment while leveraging interface-related structural characteristics that are particularly relevant for evaluating protein complex models. Together, these two complementary estimators reduce the dependence on a single accuracy estimation signal during the preliminary selection stage and facilitate the construction of a more reliable candidate subset for subsequent consensus-based ranking.

## 3 Results

### 3.1 Evaluation Metrics

Model-selection performance was evaluated using the covariance-weighted penalty adopted in CASP16 QMODE3. The component measures span global topology (TM-score), quaternary structure (QS-global, QS-best, oligo-GDTHA and oligo-GDTTS), local accuracy (lDDT), interface-local accuracy (iLDDT), and protein–protein interface quality (DockQ-ave, DockQ-wave, interface contact score (ICS) and interface patch score (IPS))[7]. Definitions of these evaluation metrics are provided in **Supplementary Note 1**. Because the component measures are correlated, the covariance-weighted penalty integrates them while limiting redundant contributions; lower values indicate that an ordered Top-5 selection is closer to the metric-specific reference quality rankings[7]. The formulation is provided in **Supplementary Note 2**.

### 3.2 Model-agnostic performance on CASP16 heteromeric and homomeric targets

To test whether DeepUMQA-Selection can identify high-quality protein complex models from large candidate pools, we retrospectively evaluated 23 CASP16 QMODE3 targets with sequence lengths ≤1,500 residues, including 14 heteromeric and 9 homomeric complexes, each represented by 8,040 candidates[7]. This experiment compares overall Top-5 penalty, target-wise performance against AlphaFold-Multimer confidence, and success rate of candidates from the high-quality region of each pool (**Figure 2**). In view of the computational demands of the two single-model estimators, which appear to scale with sequence length, we restricted their application to targets of ≤1,500 residues, a threshold within which their use remains computationally practical. For longer targets, we substituted these estimators with the intrinsic confidence metrics provided by the structure prediction models themselves (see **Section 3.5** for detailed results), thereby avoiding the prohibitive runtime that would otherwise be incurred. The subsequent USR-based alignment and consensus stages, by contrast, are comparatively lightweight and were therefore applied uniformly across all targets. This adaptive design ensures that the overall framework of the selection procedure remains intact across the full benchmark, while flexibly adapting the preliminary assessment component to target complexity without compromising computational feasibility.

**Figure 2.**
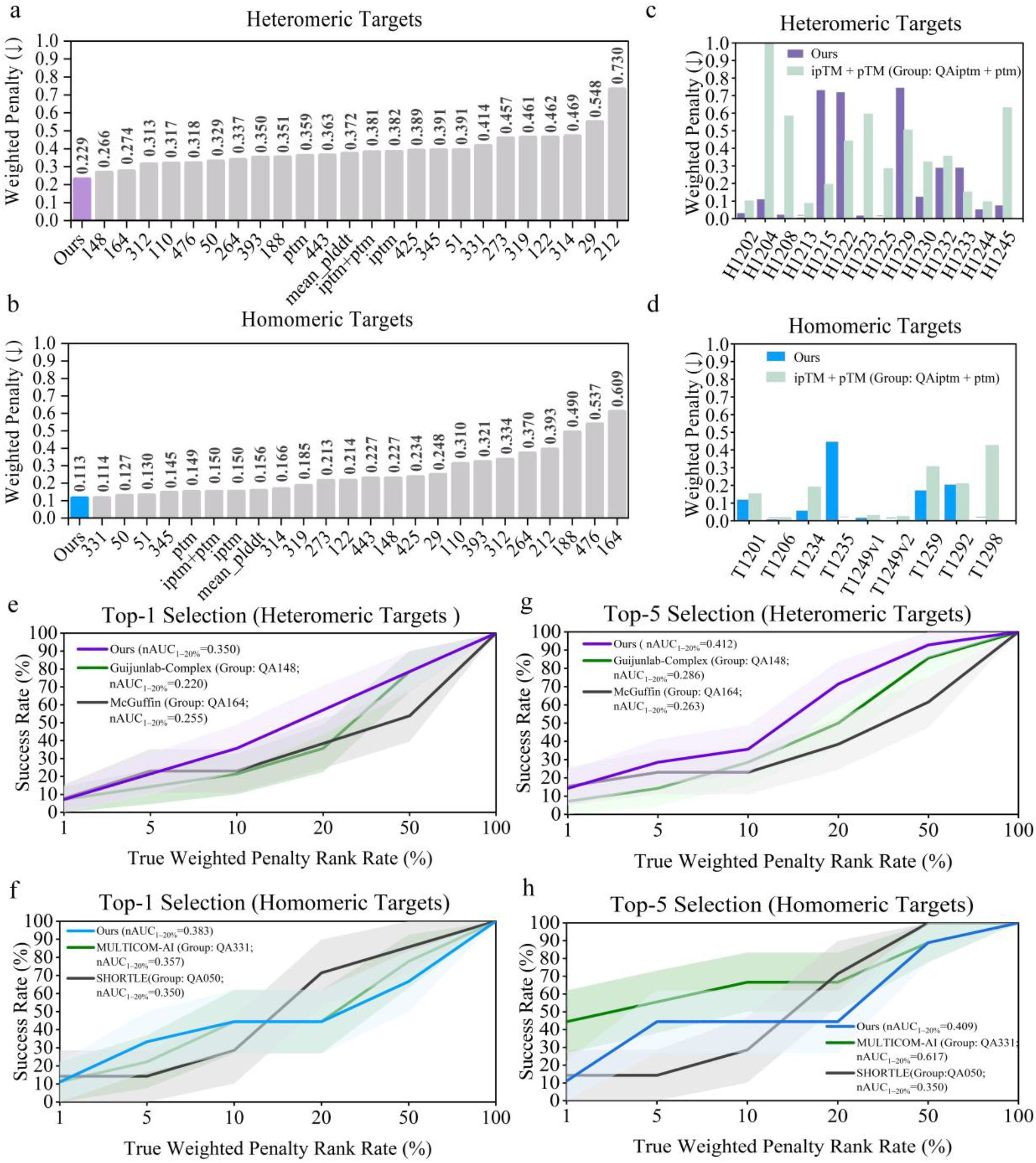
Performance of DeepUMQA-Selection on Critical Assessment of protein Structure Prediction round 16 (CASP16) targets. (a,b) Mean Top-5 weighted penalties across 14 heteromeric (a) and 9 homomeric (b) targets for DeepUMQA-Selection and the participating CASP16 groups. The component evaluation scores used to calculate the QMODE3 weighted penalty for the CASP16 participating groups included in the comparison were obtained from the official CASP16 QMODE3 evaluation data released by the CASP Prediction Center. The official QMODE3 rankings and corresponding evaluation data are available at https://predictioncenter.org/casp16/zscores_EMA.cgi and https://predictioncenter.org/download_area/CASP16/results/QA/, respectively. The y-axis shows the mean Top-5 weighted penalty, with lower values indicating better selection performance; numeric labels on the x-axis denote CASP16 group identifiers, and values above the bars indicate mean weighted penalties. (c,d) Target-wise Top-5 weighted-penalty comparisons between DeepUMQA-Selection and the predictor-derived ipTM + pTM self-assessment group (Group: QAiptm+ptm) for heteromeric and homomeric (d) targets. Target identifiers are shown on the x-axis and Top-5 weighted penalties on the y-axis; lower values indicate better performance. (e,g) Top-1 (e) and Top-5 (g) success rate for heteromeric targets, comparing DeepUMQA-Selection with Guijunlab-Complex (Group: QA148) and McGuffin (Group: QA164). (f,h) Top-1 (f) and Top-5 (h) success rate for homomeric targets, comparing DeepUMQA-Selection with MULTICOM-AI (Group: QA331) and SHORTLE (Group: QA050). For (e–h), the x-axis shows the ground-truth model-rank percentile defined by candidate-level weighted penalty, and the y-axis shows the success rate. Success rate is evaluated as the rank threshold expands from the best 1% to 100% of each 8,040-model pool; shading denotes mean ± standard error of the mean (SEM) across targets. “Ours” denotes DeepUMQA-Selection. The normalized area under the success rate curve over the 1-20% ground-truth percentile range (nAUC_1−20%_) was calculated by trapezoidal integration of the success rates at q=1,5,10, and 20%, assuming piecewise-linear interpolation between adjacent thresholds, and normalized by the 19-percentage-point interval.

We first processed antibody–antigen targets using the single-estimator, reference-guided strategy designed for general heteromeric complexes. This attempt, however, yielded unsatisfactory performance, presumably because antibody–antigen recognition is predominantly interface-driven, whereas a single global estimator that primarily evaluates overall topology is insufficient to reflect the correctness of the binding interface[36]. To overcome this limitation, we adopted the dual-estimator, hierarchical-consensus pipeline originally developed for homomeric targets. This pipeline integrates complementary assessments of global fold and interfacial quality prior to USR-based alignment, thereby furnishing a more robust and comprehensive basis for consensus ranking. Notably, although these targets were statistically grouped with heteromerics in the benchmark, they were processed using this alternative pipeline. A detailed evaluation of this tailored strategy is provided in **Sections 3.4 and 3.6**.

Selection performance was quantified by the official CASP16 QMODE3 Top-5 weighted penalty[7]. The metric measures the deviation of an ordered Top-5 model submission from metric-specific reference rankings and combines those deviations with covariance weighting; lower values indicate closer agreement with the reference quality profile. The formula for computing this weighted penalty score is given in **Supplementary Note 2**.

At the group level, DeepUMQA-Selection achieved the best mean penalty for heteromeric targets (0.229), a 13.9% reduction relative to Guijunlab-Complex (0.266; **Figure 2a**). For homomerics, its penalty (0.113) was comparable to that of MULTICOM-AI (0.114) and outperformed SHORTLE (0.127; **Figure 2b**). Thus, the aggregate gain was concentrated in heteromeric targets, whereas the homomeric result indicates parity rather than a large advantage. The numerical data underlying **Figure 2a–d** are provided in **Supplementary Table S5**.

At the target level, DeepUMQA-Selection yielded weighted penalties that outperformed the predictor-derived ipTM + pTM self-assessment group (Group: QAiptm+ptm) for 10 of 14 heteromerics (**Figure 2c**) and 8 of 9 homomerics (**Figure 2d**), or 18 of 23 targets overall. These results demonstrate that DeepUMQA-Selection offers reliable model-agnostic ranking that generalizes well across both heteromeric and homomeric targets, outperforming predictor-derived confidence in the majority of cases. Target-wise comparisons with the AlphaFold-Multimer confidence metrics ipTM, pTM, and mean pLDDT are shown in **Supplementary Figures S2 and S3**. Complete target-wise comparisons against QAiptm, QAptm and QAmean_plddt are provided in **Supplementary Tables S6 and S7**.

Success rate analysis was performed to assess whether the models selected by DeepUMQA-Selection originated from the highest-quality subset of each candidate pool (**Figure 2e–h**). A successful selection was defined as one in which the Top-1 prediction, or at least one member of the Top-5 set, fell within the lowest-penalty Top *k*% of candidates, where *k* ranges from 1 to 100. To capture success rate performance in the most demanding regime, we adopted the nAUC_1−20%_, which summarizes the success rate over the 1–20% rank-percentile interval. For heteromeric targets, DeepUMQA-Selection achieved Top-1 and Top-5 nAUC_1−20%_ values of 0.350 and 0.412 **(Figure 2e,g)**, markedly exceeding the best competing values of 0.255 and 0.286. Critically, this advantage is observed consistently at both the Top-1 and Top-5 levels, indicating that the performance gain stems from improved identification of the top-tier models in large heteromeric pools, rather than being an artifact of a lower aggregate penalty.

For homomeric targets, DeepUMQA-Selection achieved a Top-1 nAUC_1--20%_ of 0.383, compared with 0.357 for MULTICOM-AI and 0.350 for SHORTLE **(Figure 2f)**. For Top-5 success rate, DeepUMQA-Selection achieved an nAUC_1−20%_ of 0.409, which was higher than SHORTLE (0.350) but lower than MULTICOM-AI (0.617; **Figure 2h**). This contrast is more informative than the near-equal mean penalties in **Figure 2b**, because it distinguishes the ability to identify a strong first-choice model from the ability to recover at least one high-quality model within the submitted Top-5 set. The selected Top-5 models and their structural quality scores for targets with ≤1,500 residues are provided in **Supplementary Table S8**.

### 3.3 Multimetric quality assessment of DeepUMQA-Selection models

To assess the generalizability of DeepUMQA-Selection across structurally diverse protein complexes, we examined three complementary quality dimensions, namely global topology, interface quality, and local accuracy, on six representative targets (**Figure 3**): two non-antibody–antigen heteromeric complexes (H1202 and H1208), two homomeric complexes (T1206 and T1298), and two antibody– antigen complexes (H1223 and H1215). For each target, TM-score[27], DockQ-wave[9,28], and lDDT[29] were calculated over the full set of 8,040 candidate models. For each target and metric, we compared the score of the best model among all candidate models, the score of the best model among the Top-5 models selected by DeepUMQA-Selection, and the mean score of the selected Top-5 models. Representative structural overlays for H1208 and T1298 are provided in **Figure 3d,e**. It should be noted that the weighted penalty, by design, integrates multiple correlated quality metrics; consequently, the model that minimizes this penalty may not be the best performer on any single metric, but rather represents an optimal trade-off across all evaluated aspects of structural quality.

**Figure 3.**
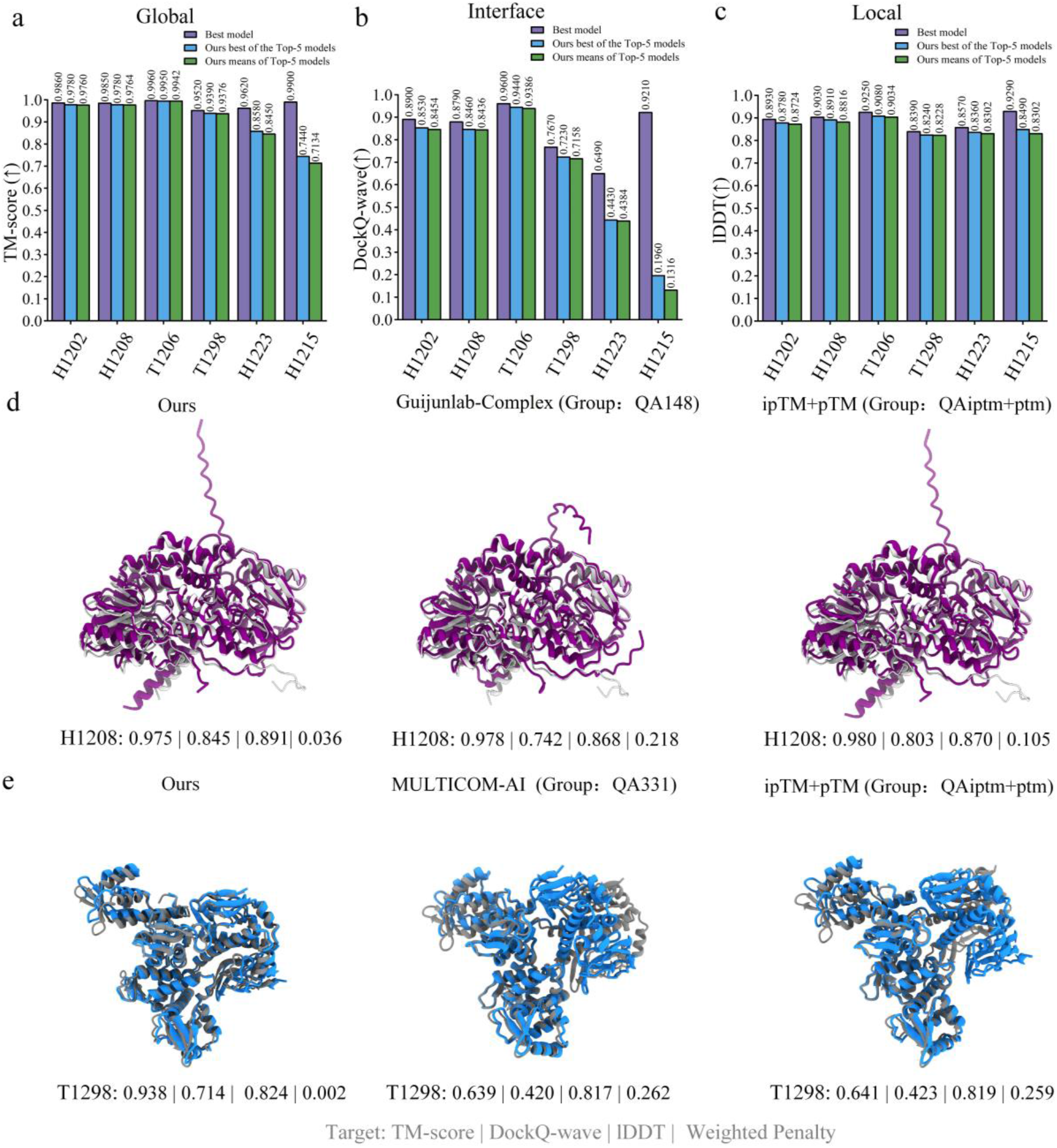
Multimetric quality of selected models. (a–c) Comparison of TM-score (global topology), DockQ-wave (interface quality), and lDDT (local accuracy) for H1202, H1208, T1206, T1298, H1223, and H1215. For each target and metric, the purple bars show the score of the best model among all 8,040 candidate models, the blue bars show the score of the best model among the Top-5 models selected by DeepUMQA-Selection, and the green bars show the mean score of the selected Top-5 models. (d,e) Structural superpositions compare the H1208 (d) and T1298 (e) models selected by DeepUMQA-Selection and the indicated comparison methods. Grey structures denote the experimental references, whereas the selected models are shown in purple for H1208 (d) and blue for T1298 (e). Values below each structure are reported as TM-score | DockQ-wave | lDDT | Weighted Penalty. Higher TM-score, DockQ-wave, and lDDT values and lower weighted penalties indicate better performance.

For the non-antibody–antigen heteromeric and homomeric targets, the scores of the best models among the selected Top-5 were generally close to those of the best models among all 8,040 candidate models across TM-score, DockQ-wave, and lDDT (**Figure 3a–c**). The Top-5 mean scores were also close to the corresponding best scores among the selected Top-5 for most targets, indicating that favorable structural quality was maintained across multiple selected models rather than being confined to a single representative. Antibody–antigen complexes, however, constituted a notable exception, with H1215 being a particularly instructive case: the selected models achieved lDDT values approaching 0.8, yet showed substantially lower DockQ-wave scores and only moderate TM-scores. This observation highlights that local structural plausibility does not guarantee correct interchain assembly, a limitation that would remain undetected by global or local quality scores alone.

Comparison of the representative structures pinpoints the specific quality dimension that accounts for the observed ranking differences. In the case of H1208 (**Figure 3d**), all three methods delivered comparable TM-scores (0.975–0.980), yet DeepUMQA-Selection achieved superior DockQ-wave (0.845) and lDDT (0.891) values alongside the best weighted penalty (0.036), indicating that its advantage originated from interface and local accuracy rather than global topology. For T1298 (**Figure 3e**), DeepUMQA-Selection attained TM-score, DockQ-wave, and lDDT values of 0.938, 0.714, and 0.824, respectively, while the competing methods showed similar lDDT but markedly inferior global and interface performance. This demonstrates that the multimetric penalty distinguished models that would have appeared nearly identical when assessed by local accuracy alone.

### 3.4 Performance on alternative conformational states and antibody–antigen targets

We further evaluated DeepUMQA-Selection in two challenging model-selection scenarios: alternative conformational states and antibody–antigen complexes (**Figure 4**). Protein conformational changes are closely linked to biological function, and the same complex may adopt distinct functionally relevant states under different conditions, posing a challenge for model selection when candidate models span multiple conformational states. Antibody–antigen complexes pose another challenge for model selection because their highly specific recognition is primarily mediated by the complementarity-determining regions (CDRs), particularly the structurally diverse and conformationally flexible CDR-H3 loop, resulting in substantial variation in interface accuracy among candidate models and complicating the identification of high-quality complexes. In CASP16, two protein-complex targets were associated with experimentally resolved alternative conformational states, T1249 and T1294[37]. We evaluated DeepUMQA-Selection on T1249, which is a homotrimeric spike protein from Sabiá virus that adopts a compact closed state (T1249v1, PDB ID: 9FYA) and, under acidic conditions and upon binding an unidentified metal ion, an open state (T1249v2, PDB ID: 9FYG)[37], shown on the left and right of **Figure 4a**, respectively. T1294 was excluded from the detailed CASP16 assessment because its two experimentally resolved states showed only limited structural differences[37]. For the antibody–antigen complexes, we evaluated DeepUMQA-Selection on the eight antibody–antigen targets in the CASP16 QMODE3 benchmark dataset.

**Figure 4.**
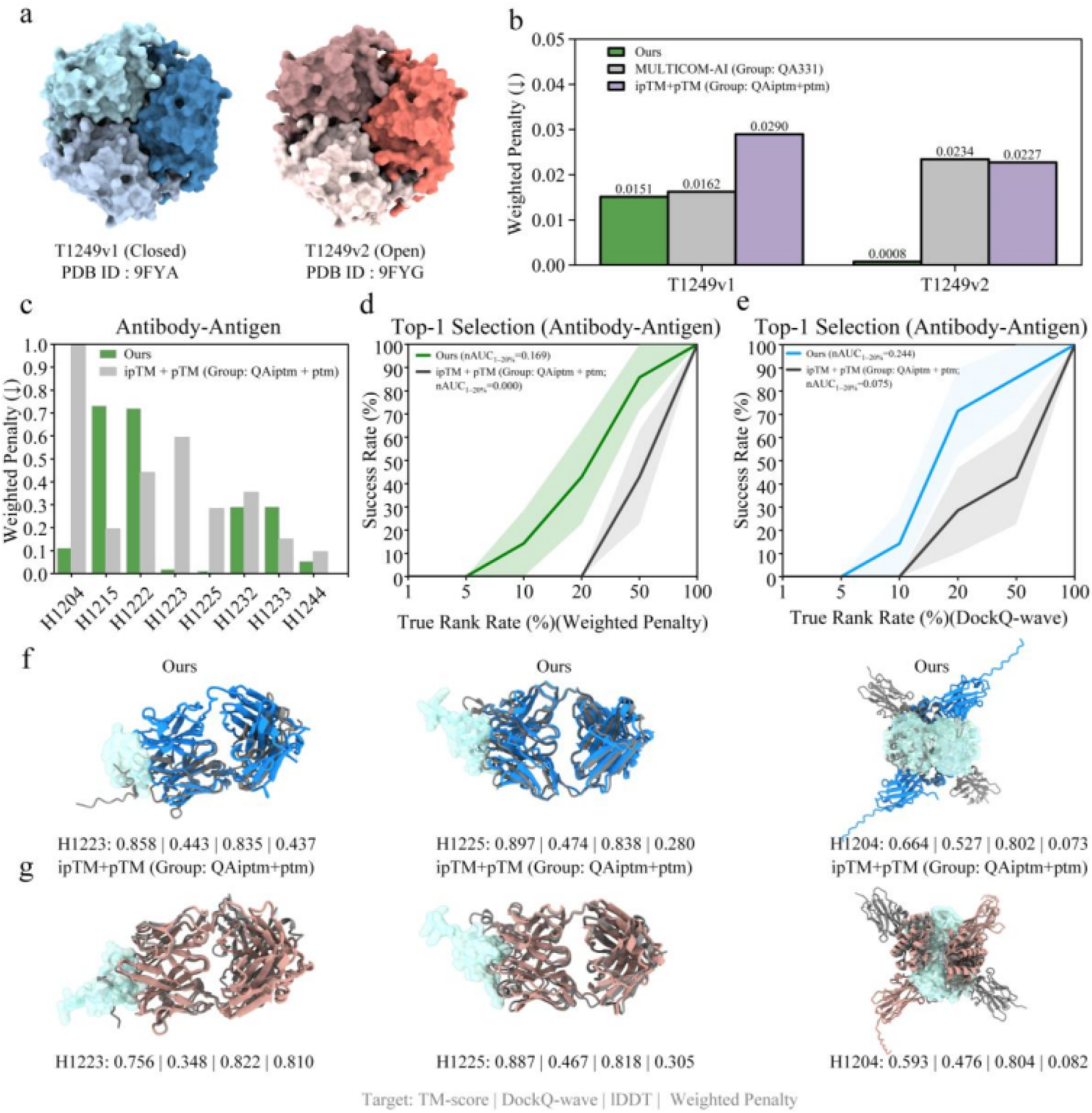
State-specific T1249 selection and antibody–antigen evaluation. (a) Reference structures of the closed T1249v1 (Protein Data Bank ID: 9FYA) and open T1249v2 (9FYG) conformational states. (b) Top-5 weighted penalties obtained by DeepUMQA-Selection (Ours), MULTICOM-AI (Group: QA331), and the predictor-derived ipTM+pTM self-assessment baseline (Group: QAiptm+ptm) against the two reference states. (c) Target-wise Top-5 weighted penalties for eight antibody–antigen targets, comparing DeepUMQA-Selection with ipTM+pTM. (d,e) Top-1 model-selection success rates across seven antibody–antigen targets after excluding H1222, using weighted penalty (d) and DockQ-wave (e) as ground-truth ranking criteria. nAUC_1−20%_ values for the seven-target analysis are reported in the legends. The recovery curves for all eight antibody–antigen targets are provided in **Supplementary Figure S4**. nAUC_1−20%_ values are reported in the legends, and shading denotes mean ± standard error across targets. (f,g) Structural comparison of representative models for the antibody– antigen targets H1223, H1225, and H1204. For each target, the model with the best weighted penalty among the Top-5 models selected by DeepUMQA-Selection is shown in (f), whereas the model with the best weighted penalty among the Top-5 models ranked by the predictor-derived confidence metric ipTM+pTM (Group: QAiptm+ptm) is shown in (g). The experimental reference complex is shown in grey, while the antibody/nanobody and antigen components of the selected models are shown in blue and green for DeepUMQA-Selection and in salmon and cyan for ipTM+pTM, respectively. Higher TM-score, DockQ-wave, and lDDT values indicate higher structural accuracy, whereas lower weighted penalties indicate more accurate model selection.

For T1249, DeepUMQA-Selection selected Top-5 models from the common candidate pool, which was then evaluated separately against the experimentally resolved closed (T1249v1) and open (T1249v2) conformational states (**Figure 4a,b**). We compared DeepUMQA-Selection with the second-ranked method MULTICOM-AI (Group: QA331) and the predictor-derived confidence score ipTM+pTM (Group: QAiptm+ptm) using the weighted penalty (**Figure 4b**). For the closed state T1249v1, DeepUMQA-Selection achieved the best selection performance, with a weighted penalty of 0.0151, slightly outperforming MULTICOM-AI (0.0162) and substantially outperforming ipTM+pTM (0.0290). For the open state T1249v2, the advantage was more pronounced, with DeepUMQA-Selection achieving a weighted penalty of 0.0008, compared with 0.0234 for MULTICOM-AI and 0.0227 for ipTM+pTM. These results show that DeepUMQA-Selection achieved favorable selection performance for both conformational states, with a more pronounced advantage for the open state. The consistent advantage observed against both T1249v1 and T1249v2 indicate that DeepUMQA-Selection maintained high selection quality when evaluated against either experimentally resolved state, highlighting the potential of DeepUMQA-Selection for model selection in protein complexes with alternative conformational states.

For the eight antibody–antigen targets, DeepUMQA-Selection outperformed predictor-derived confidence metric ipTM+pTM on five targets according to the Top-5 weighted penalty, whereas ipTM+pTM performed better for the three targets (**Figure 4c**). Among these targets, we found that H1222 appeared as an outlier. To reduce the potential statistical bias introduced by this target, we excluded H1222 and performed the success rate analysis on the remaining seven targets (**Figure 4d,e**). When candidate models were ranked by weighted penalty, DeepUMQA-Selection achieved a higher Top-1 nAUC_1−20%_ of 0.169 than ipTM+pTM (0.000) (**Figure 4d**). When candidate models were ranked by DockQ-wave, DeepUMQA-Selection also achieved a higher Top-1 nAUC_1−20%_ of 0.244 than ipTM+pTM (0.075) (**Figure 4e**). The superior Top-1 nAUC_1−20%_ values achieved by DeepUMQA-Selection under both ranking criteria indicate a stronger ability to recover high-quality models within the top-ranked region of the candidate pool. The consistent advantage observed for both weighted penalty and DockQ-wave further demonstrates that DeepUMQA-Selection not only improves overall model ranking but also enhances the recovery of candidates with accurate antibody–antigen interfaces, supporting its effectiveness for high-quality model selection in antibody–antigen complexes. The corresponding Top-1 success rate analysis including all eight antibody–antigen targets are provided in **Supplementary Figure S4**.

We selected three representative targets, H1223, H1225, and H1204, to compare the quality of the best models from the respective Top-5 models selected by DeepUMQA-Selection (**Figure 4f**) and the predictor-derived confidence score ipTM+pTM (**Figure 4g**). For H1223, DeepUMQA-Selection showed the most pronounced improvement, achieving a weighted penalty of 0.437 compared with 0.810 for ipTM+pTM, corresponding to a 46.0% improvement, while increasing TM-score from 0.756 to 0.858 and DockQ-wave from 0.348 to 0.443, increases of 13.5% and 27.3%, respectively. For H1225, DeepUMQA-Selection showed more modest but consistent improvement, achieving a weighted penalty of 0.280 compared with 0.305 for ipTM+pTM, corresponding to an 8.2% improvement. For H1204, DeepUMQA-Selection achieved a weighted penalty of 0.073, compared with 0.082 for ipTM+pTM, representing an 11.0% improvement, while increasing TM-score from 0.593 to 0.664 and DockQ-wave from 0.476 to 0.527, increases of 12.0% and 10.7%, respectively. Across these representative cases, models with more favorable weighted penalty values generally also achieved higher scores on individual structural quality metrics, indicating that improved performance according to the integrated selection criterion was typically accompanied by corresponding gains in global topology, interface, and local structural accuracy. This consistency across complementary quality measures is consistent with the weighted penalty used in CASP16 QMODE3 to integrate multiple aspects of model quality while accounting for correlations among component metrics.

### 3.5 Model selection performance on the all CASP16 complex set

To evaluate the scalability of DeepUMQA-Selection across protein complexes with varying sequence lengths, we introduced a confidence-assisted strategy for large targets, in which predictor-derived confidence metrics were incorporated into the preliminary selection stage. For large targets with sequence lengths >1,500 residues, predictor-derived confidence metrics were used in place of the model-agnostic single-model quality estimators. Specifically, for large heteromeric targets, EMA_SIT_ was replaced by AlphaFold-Multimer ranking confidence (0.8×ipTM+0.2×pTM)[25,26], whereas for large homomeric targets, EMA_SGT_ and EMA_SIT_ were replaced by mean pLDDT and AlphaFold-Multimer ranking confidence, respectively. All subsequent branch-specific consensus-selection procedures were applied without modification. The detailed confidence-assisted workflow for large targets is shown in **Supplementary Figure S5**. For targets with ≤1,500 residues, the standard DeepUMQA-Selection workflow was retained without modification. The target lists for the standard and confidence-assisted workflows are provided in **Supplementary Tables S2 and S3**, respectively. We then evaluated the model-selection performance of DeepUMQA-Selection across all 36 CASP16 QMODE3 protein complex targets (**Figure 5**).

**Figure 5.**
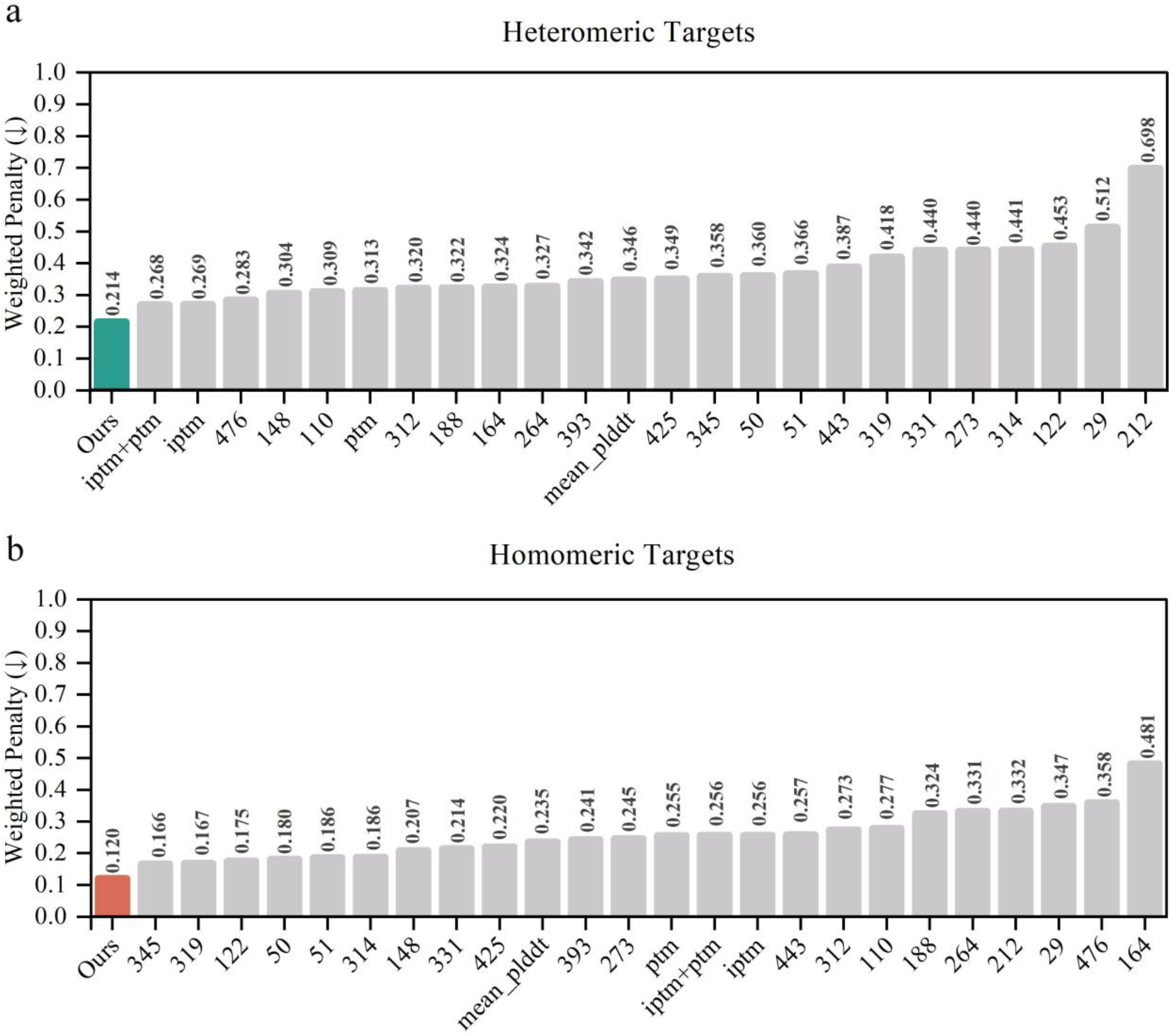
Performance on the complete CASP16 QMODE3 protein complex benchmark. (a,b) Mean Top-5 weighted penalties are shown for (a) 21 heteromeric and (b) 15 homomeric targets; lower weighted penalty scores indicate better model selection performance. Numeric x-axis labels denote CASP16 group identifiers, predictor-derived confidence baselines are labeled by metric name, and “Ours” denotes DeepUMQA-Selection.

For the 21 heteromeric targets, we compared DeepUMQA-Selection with both predictor-derived confidence metrics and participating CASP16 EMA methods (**Figure 5a**). Among the predictor-derived confidence metrics, ipTM+pTM showed the best model-selection performance for heteromeric targets in CASP16, with the lowest weighted penalty of 0.268. DeepUMQA-Selection achieved a weighted penalty of 0.214, compared with 0.268 for ipTM+pTM, representing a 20.1% improvement. It also outperformed ipTM (0.269), pTM (0.313), and mean pLDDT (0.346), with improvements of 20.4%, 31.6%, and 38.2%, respectively. Among the participating CASP16 EMA groups excluding predictor-derived self-assessment methods, VifChartreuse (Group: QA476) achieved the lowest weighted penalty of 0.283. DeepUMQA-Selection achieved a weighted penalty of 0.214, representing a 24.4% improvement over VifChartreuse. These results indicate that DeepUMQA-Selection improves model-selection performance for heteromeric complexes, while the relatively high weighted penalties observed across methods underscore the substantial room for further improvement.

For the 15 homomeric targets (**Figure 5b**), model-selection performance was generally better than for heteromeric complexes, as reflected by the weighted penalty. DeepUMQA-Selection outperformed all predictor-derived confidence baselines, achieving a weighted penalty of 0.120, compared with 0.235 for mean pLDDT, 0.255 for pTM, 0.256 for ipTM, and 0.256 for ipTM+pTM, representing improvements ranging from 48.9% to 53.1% over these baselines. It also outperformed MULTICOM_human (Group: QA345), the best-performing participating CASP16 group, achieving a weighted penalty of 0.120 compared with 0.166 for MULTICOM_human, representing a 27.7% improvement. The selected Top-5 candidate models and their weighted penalties and evaluation metric values for CASP16 homomeric and heteromeric targets with sequence lengths >1,500 residues are provided in **Supplementary Table S9**.

These results support the scalability of the DeepUMQA-Selection pipeline, demonstrating that its preliminary assessment stage can be adapted for large targets while preserving the downstream consensus-selection framework and maintaining effective model-selection performance. The superior model-selection performance of DeepUMQA-Selection relative to direct confidence-based ranking, as reflected by its more favorable weighted penalty, further suggests that the downstream consensus-selection procedure may provide additional discriminatory power beyond predictor-derived confidence alone, thereby contributing to more effective identification of high-quality models from large candidate pools.

### 3.6 Ablation analysis and computational efficiency of USR-based structural alignment

To evaluate the contributions of individual components to the model-selection performance of DeepUMQA-Selection and the computational efficiency of USR-based structural alignment (USR-align), we used the 23 CASP16 QMODE3 targets with sequence lengths ≤1,500 residues to perform ablation analysis for the homomeric/antibody–antigen workflow, reference set size parameter analysis for the heteromeric workflow, and runtime benchmarking of USR-align against US-align (**Figure 6**). For the homomeric/antibody–antigen workflow, we evaluated variants without iterative refinement (w/o IR) or random grouping (w/o RG), as well as direct Top-5 model selection based solely on EMA_SGT_ or EMA_SIT_ rankings (**Figure 6a**). For the non-antibody–antigen heteromeric workflow, we examined the effect of the reference set size parameter (*N*) on model-selection performance (**Figure 6b**). We compared the runtime of USR-align with that of US-align across different candidate pool sizes, with runtimes averaged across all targets at each pool size (**Figure 6c**). We further compared their runtimes across different target sequence lengths using 100 randomly sampled models per target (**Figure 6d**). All runtime benchmarks were performed under the same hardware configuration, as detailed in **Supplementary Table S10**.

**Figure 6.**
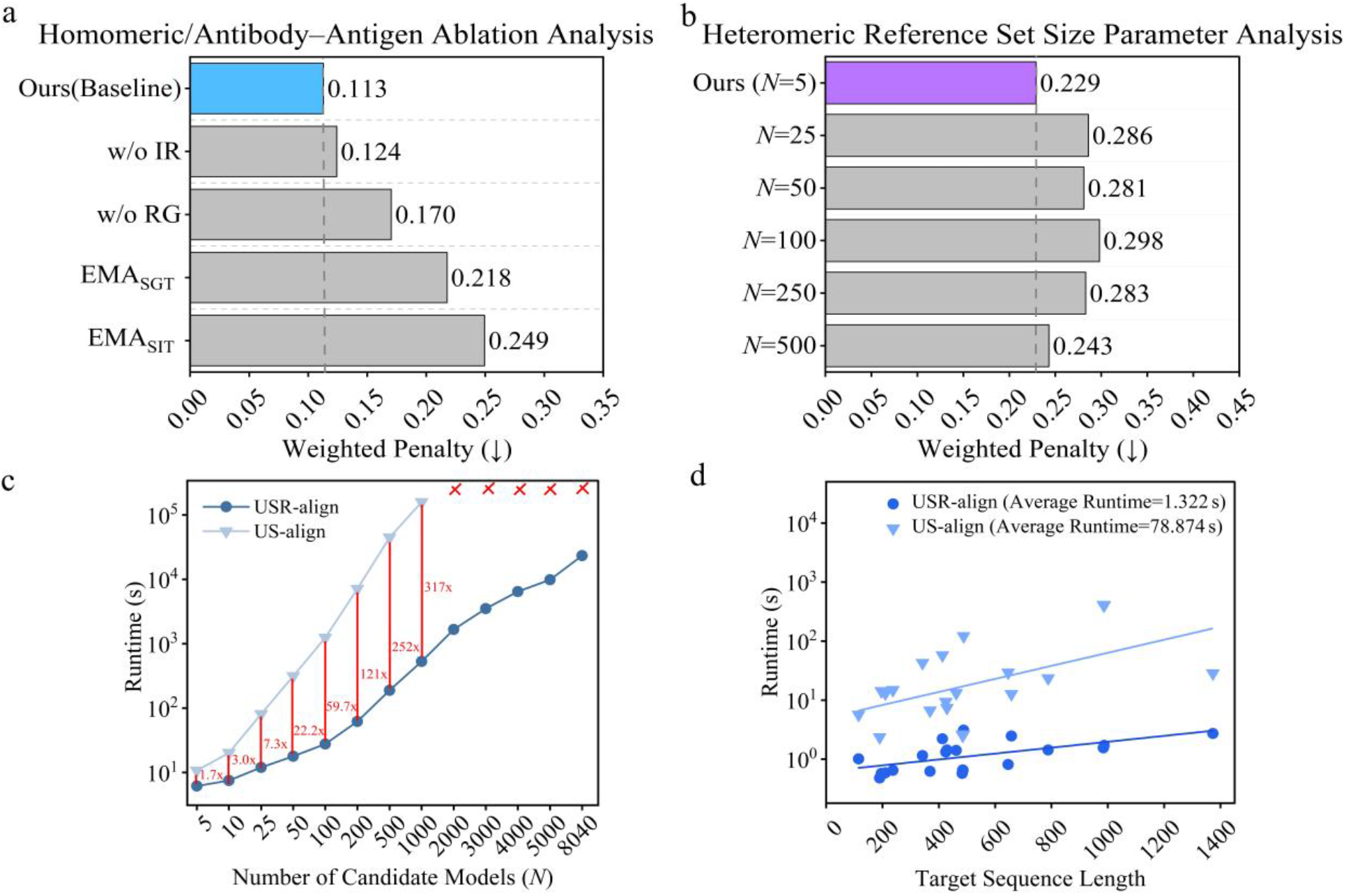
Ablation analysis and computational efficiency of USR-based structural alignment (USR-align). The analyses were based on the 23 CASP16 QMODE3 targets with sequence lengths ≤1,500 residues, with workflow-specific subsets used for (a) and (b). (a) For the homomeric/antibody–antigen workflow, mean Top-5 weighted penalties are shown for the complete DeepUMQA-Selection workflow (Ours), its ablated variants without iterative refinement (w/o IR) or random grouping (w/o RG), and direct Top-5 model selection based solely on EMA_SGT_ or EMA_SIT_ rankings; lower values indicate better model-selection performance. (b) Effect of the reference set size parameter (*N*) on model-selection performance for non-antibody–antigen heteromeric complexes; the default value was set to *N* = 5. Dashed lines in (a,b) indicate the mean Top-5 weighted penalty achieved by the complete DeepUMQA-Selection workflow, which serves as the baseline for comparison. (c) Runtime comparison of USR-align and US-align across different candidate pool sizes, with runtimes averaged across all 23 targets at each pool size. Red labels indicate the corresponding speedups of USR-align over US-align for each candidate pool sizes (*N*), and red crosses indicate US-align runs that did not complete. (d) comparison of USR-align and US-align across different target sequence lengths. For each target, 100 models were randomly sampled from the candidate pool (*N*=100) for runtime evaluation. The y-axis uses a logarithmic scale. Solid lines show log-linear regression fits. The mean runtimes across the 23 targets were 1.322 s for USR-align and 78.874 s for US-align, respectively.

For the homomeric/antibody–antigen workflow, the complete DeepUMQA-Selection framework (Ours) achieved a mean Top-5 weighted penalty of 0.113, which served as the baseline (**Figure 6a**). Removing IR increased the mean weighted penalty from 0.113 to 0.124 (a 9.7% relative increase), indicating a modest reduction in model-selection performance. The modest performance reduction suggests that IR provides an incremental benefit beyond the preliminary selection stage, which may already enrich the candidate pool with high-quality models. In contrast, removing RG resulted in a substantially greater reduction in model-selection performance, with the mean weighted penalty increasing from 0.113 to 0.170 (a 50.4% relative increase). This result highlights the effectiveness of random grouping in the hierarchical consensus selection process, which may reduce the influence of overrepresented structural conformations and facilitate the retention of high-quality candidates for subsequent consensus evaluation. A more pronounced reduction in model-selection performance was observed when the Top-5 models were selected based solely on EMA_SGT_ or EMA_SIT_ rankings, with mean weighted penalties increasing from 0.113 for the baseline to 0.218 and 0.249, respectively, representing increases of 92.9% and 120.4%. These results suggest that the combined use of complementary single-model accuracy estimates and structural consensus information improves model-selection performance compared with selection based on either estimator alone.

For non-antibody–antigen heteromeric complexes, we further examined the effect of the reference set size parameter (*N*) on model-selection performance (**Figure 6b**). The default setting of *N* = 5 served as the baseline and achieved the best model-selection performance among the tested settings, with a mean Top-5 weighted penalty of 0.229. When the reference set size was increased to *N* = 25, 50, 100, and 250, the mean Top-5 weighted penalties were 0.286, 0.281, 0.298, and 0.283, respectively. Although the differences among these settings were relatively small, all four settings resulted in a marked reduction in model-selection performance relative to the baseline, with mean weighted penalties increasing by 22.7%–30.1%. Further increasing the reference set size to *N* = 500 reduced the mean Top-5 weighted penalty to 0.243, indicating improved model-selection performance compared with the *N* = 25–250 settings, although it remained inferior to the baseline (*N* = 5). These results suggest that the preliminary single-model assessment may already identify a small set of high-quality reference models sufficient to support effective consensus-based selection. Expanding the reference set did not consistently improve model-selection performance, possibly because the inclusion of additional lower-quality reference models may introduce noise into the consensus signal. Larger reference sets also increase the computational cost of structural comparison. We therefore selected *N* = 5 as the default reference set size, balancing model-selection performance and computational efficiency.

We further evaluated the computational efficiency of USR-align by comparing its runtime with that of US-align across different candidate pool sizes under the same hardware configuration (**Figure 6c**). We observed that for *N*=10, 100, and 1,000, USR-align required 0.36, 1.32, and 21.80 s, respectively, whereas US-align required 1.10, 78.90, and 6,915.00 s, representing speedups of 3.0-, 59.7-, and 317-fold, respectively. The computational advantage of USR-align became increasingly pronounced as the candidate pool size increased. For candidate pools containing more than 1,000 models, US-align did not succeed under the tested hardware configuration, whereas USR-align remained computationally feasible for pools containing up to 8,040 models. These results demonstrate the computational advantage of USR-align for large-scale candidate pools, supporting its potential as a coarse-grained structural comparison strategy in consensus-based model quality assessment. We further examined the effect of target sequence length on the runtime of both USR-align and US-align (**Figure 6d**). For each target with the sequence length ≤1,500 residues, 100 models were randomly sampled from the candidate pool, and both methods were evaluated using the same sampled set (*N* = 100). We observed that USR-align consistently exhibited substantially shorter runtimes than US-align across the evaluated targets, with mean runtimes of 1.322 and 78.874 s, respectively, yielding a speedup of approximately 59.7-fold based on the mean runtimes. Although the runtime of both methods generally increased with target sequence length, USR-align exhibited a more gradual increase in runtime and consistently required substantially less time than US-align across the evaluated targets, demonstrating its computational efficiency over the evaluated range of target sequence lengths.

## 4 Conclusion

In this study, we developed DeepUMQA-Selection, a two-stage framework for selecting high-quality protein complex models from large-scale candidate pools. By first identifying a quality-enriched subset of candidate models through single-model accuracy estimators and subsequently applying USR-based structural alignment for consensus ranking, our method reduces the number of structural comparisons while avoiding computationally expensive structural superposition. On the retrospective CASP16 QMODE3 benchmark, DeepUMQA-Selection outperformed all participating methods and predictor-derived confidence metrics for targets with sequence lengths ≤1,500 residues, with particularly pronounced improvements for heteromeric targets, whose structurally heterogeneous candidate pools pose particular challenges for reliable model selection. The confidence-assisted extension further enabled the framework to accommodate larger targets. Across the complete 36 targets, DeepUMQA-Selection outperformed all competing methods, demonstrating the scalability of the two-stage framework and the effectiveness of integrating predictor-derived confidence metrics with USR-based structural consensus. Ablation analyses further supported the contributions of the key components to overall model-selection performance, while runtime benchmarking demonstrated the substantial computational advantage of USR-based structural comparison over US-align, with a 317-fold speedup for candidate pools containing 1,000 models. These results suggest DeepUMQA-Selection as an accurate and computationally efficient framework for selecting high-quality protein complex models from large-scale candidate pools.

Building on these findings, future work could further improve the accuracy and computational efficiency of the single-model accuracy estimators, with greater computational efficiency enabling the fully model-agnostic workflow to be extended to larger protein complexes. Another direction is to explore the integration of predictor-derived confidence metrics with independent accuracy estimates to improve preliminary model selection. Given that protein complexes may adopt multiple biologically relevant conformations, DeepUMQA-Selection could be further extended to identify high-quality models representing distinct conformational states. In addition, incorporating structural and physicochemical information, such as all-atom structural validation metrics[38] and energy-based descriptors[39], could complement existing model-selection criteria by providing quality information on local stereochemical quality or interchain interactions.

## Key points

- We develop DeepUMQA-Selection, an accurate and efficient framework for large-scale protein complex model selection, with a model-agnostic standard workflow and a confidence-assisted extension for large targets.
- A two-stage strategy combining complementary single-model assessment and USR-based structural consensus enables scalable model prioritization.
- DeepUMQA-Selection achieved the best performance in the retrospective CASP16 QMODE3 benchmark.
- USR-based structural similarity evaluation reduces computational cost by approximately 317-fold compared with exhaustive pairwise US-align comparisons.

## Data availability

The CASP16 QMODE3 benchmark dataset used in this study is publicly available through the CASP Prediction Center (https://predictioncenter.org/download_area/CASP16/results/QA/). Benchmark evaluation scripts (https://github.com/alisiafadini/casp16/blob/main/QMODE3/). The processed candidate model sets, and evaluation results generated in this study can be made available at https://github.com/GBLille/CASP16-CAPRI_MassiveFold_Data.

## Web server availability

The DeepUMQA-Selection web server will be made freely available upon public release.

## Funding

This work was supported by the National Key R&D Program of China [2022ZD0115103, G.Z.], the National Nature Science Foundation of China [62573386, G.Z.], the “Pioneer” and “Leading Goose” R&D Program of Zhejiang [2025C01190, G.Z.], and the Zhejiang Province High-level Talent Special Support Program [2023R5248, G.Z.].

## Acknowledgements

We thank the College of Information Engineering at Zhejiang University of Technology for providing the computational resources used in this study.

## Author contributions

Guijun Zhang conceived and supervised the research. Guijun Zhang, Fang Liang and Lei Xie designed the experiments. Fang Liang, Lei Xie, Enjia Ye, Zhengri Pan, Tiantian Xu, Haodong Wang, and Pengcheng Wei collected the data and performed the experiments. Guijun Zhang, Fang Liang and Lei Xie analyzed the data. Guijun Zhang, Fang Liang and Lei Xie wrote the manuscript. Guijun Zhang and Jun Liu revised the manuscript. All authors read and approved the final manuscript.

## Conflict of interest

The authors declare no competing interests.

## Supplementary Information

### Supplementary Figures

**Supplementary Figure S1.**
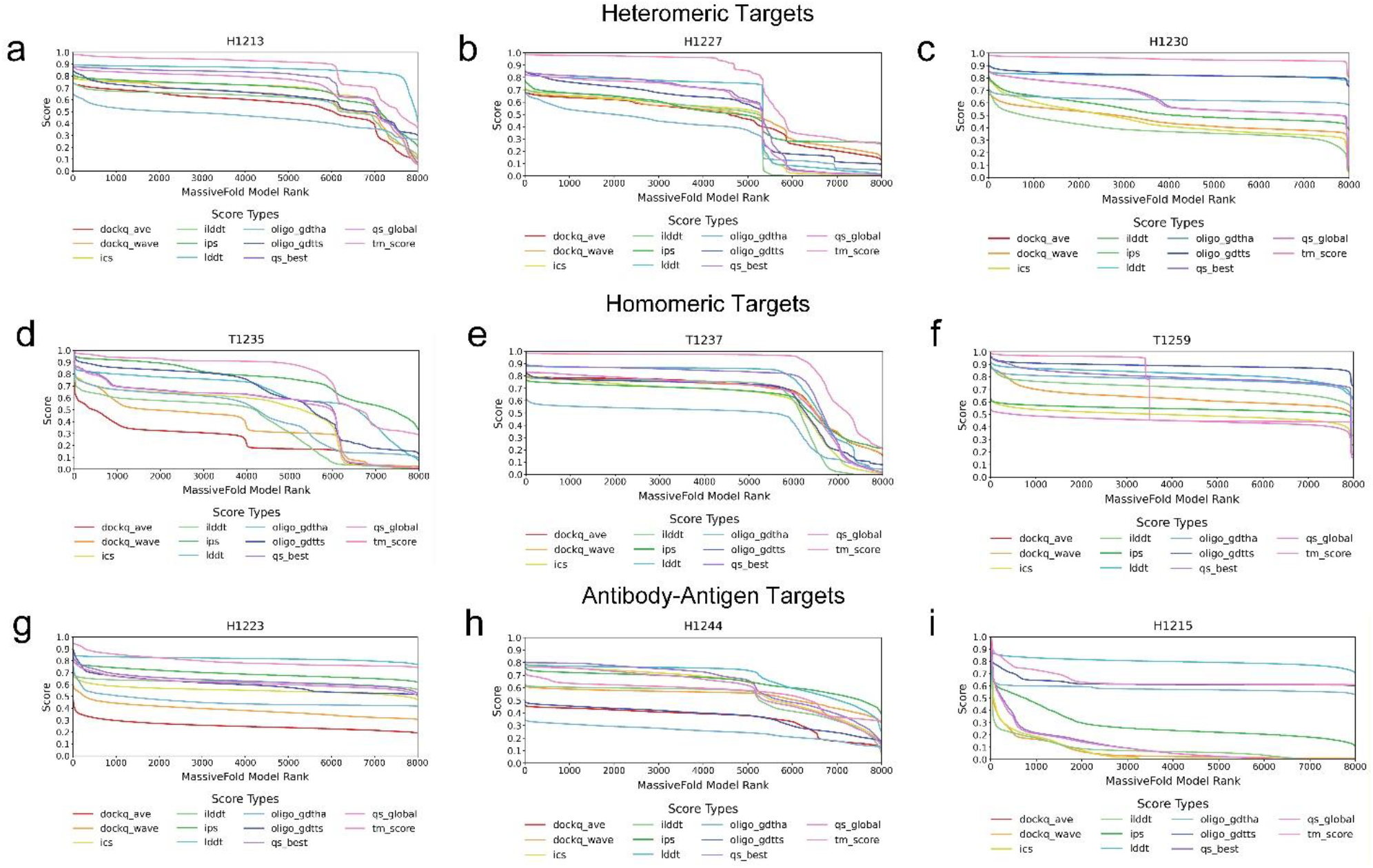
Quality distributions of MassiveFold models across representative targets. Each MassiveFold model for three heteromeric targets (H1213 (a), H1227 (b), and H1230 (c)), three homomeric targets (T1235 (d), T1237 (e), and T1259 (f)), and three antibody–antigen targets (H1223 (g), H1244 (h), and H1215 (i)) was compared with the corresponding experimental structure using the 11 evaluation metrics included in the weighted penalty as those visualized in the CASP16 QMODE3 assessment. For each metric, the candidate models were ranked in descending order according to their scores, and the metric score was plotted against the resulting metric-specific model rank. These metrics were selected to facilitate direct comparison across different aspects of structural quality, as all have values ranging from 0 to 1, with higher scores indicating higher structural accuracy. RMSD was excluded from the visualization because it is measured on a different scale and lower values indicate higher structural accuracy. (The data were obtained from https://github.com/alisiafadini/casp16/tree/main/QMODE3/data/per_score_rankings.)

**Supplementary Figure S2.**
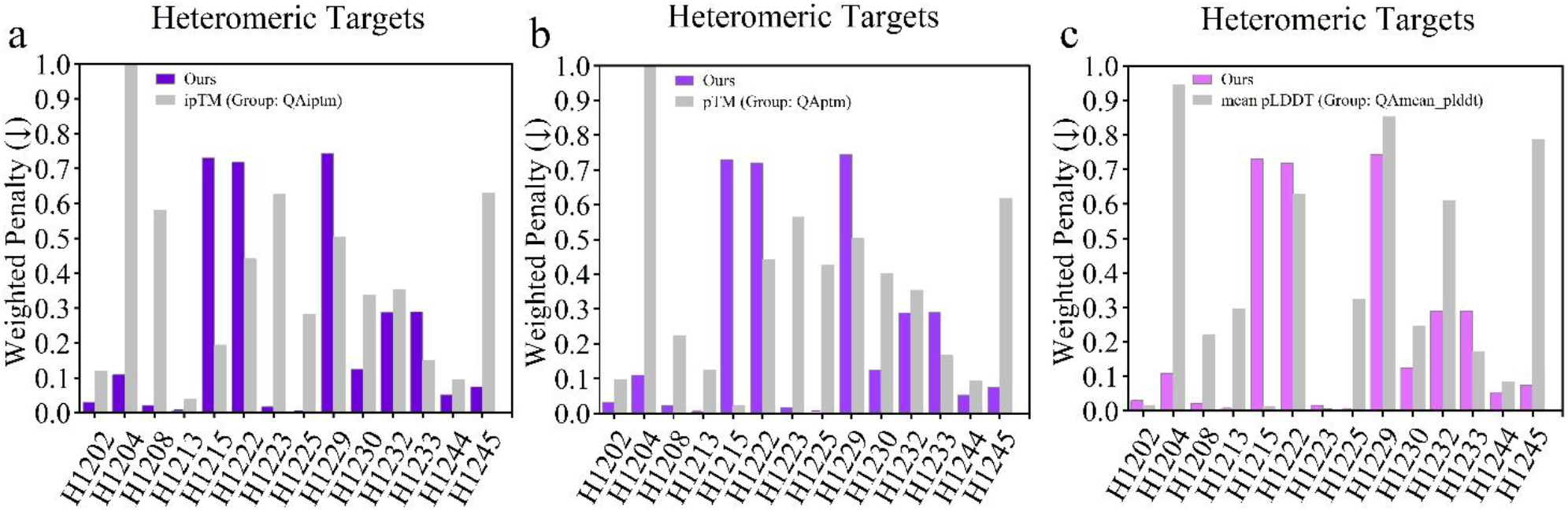
Target-wise performance of DeepUMQA-Selection and CASP16 predictor-confidence groups on heteromeric targets. (a–c) Weighted-penalty comparisons between DeepUMQA-Selection and the CASP16 groups QAiptm (a), QAptm (b), and QAmean_plddt (c) for 14 heteromeric targets. Lower values indicate better model-selection performance. The x-axis labels denote CASP16 target identifiers. “Ours” denotes DeepUMQA-Selection.

**Supplementary Figure S3.**
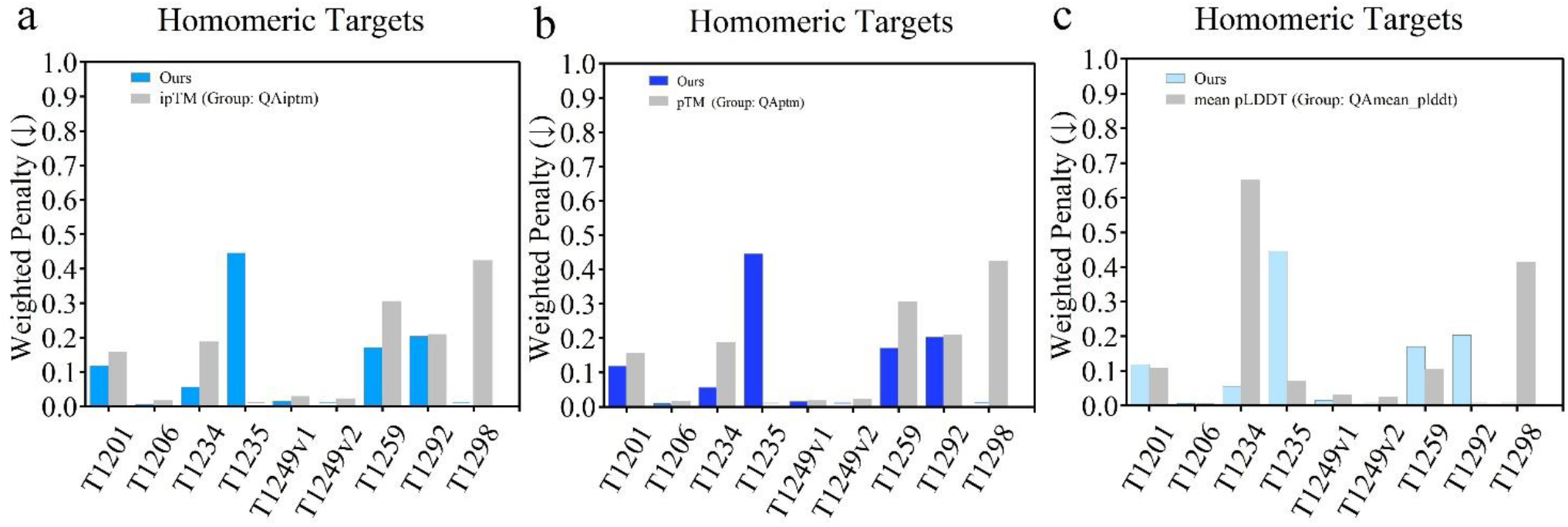
Target-wise performance of DeepUMQA-Selection and CASP16 predictor-confidence groups on homomeric targets. (a–c) Weighted-penalty comparisons between DeepUMQA-Selection and the CASP16 groups QAiptm (a), QAptm (b), and QAmean_plddt (c) for 9 homomeric targets. Lower values indicate better model-selection performance. The x-axis labels denote CASP16 target identifiers. “Ours” denotes DeepUMQA-Selection.

**Supplementary Figure S4.**
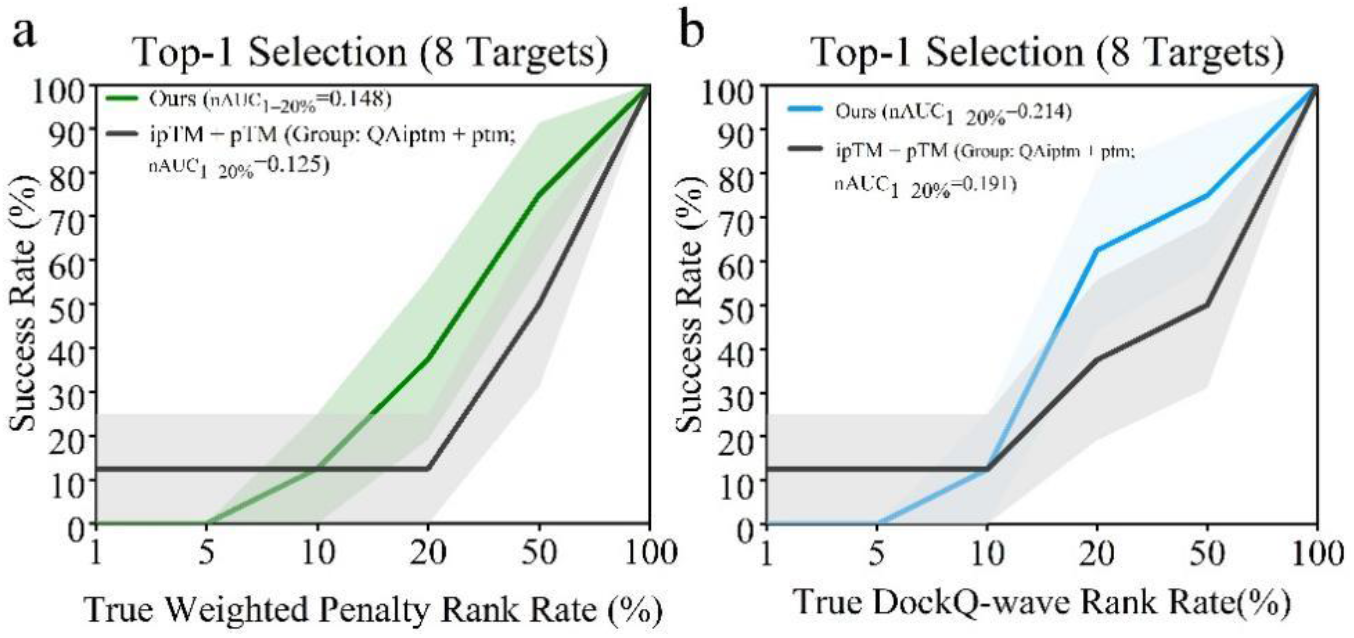
Top-1 success rate analysis for eight antibody–antigen targets. (a,b) Top-1 model-selection success rates across all eight antibody–antigen targets, using weighted penalty (a) and DockQ-wave (b) as ground-truth ranking criteria. The nAUC_1−20%_ values for the eight-target analysis are reported in the legends. For the weighted penalty analysis, the nAUC_1−20%_ values were 0.148 and 0.125 for DeepUMQA-Selection (Ours) and ipTM+pTM, respectively. For the DockQ-wave analysis, the corresponding values were 0.214 and 0.191, respectively.

**Supplementary Figure S5.**
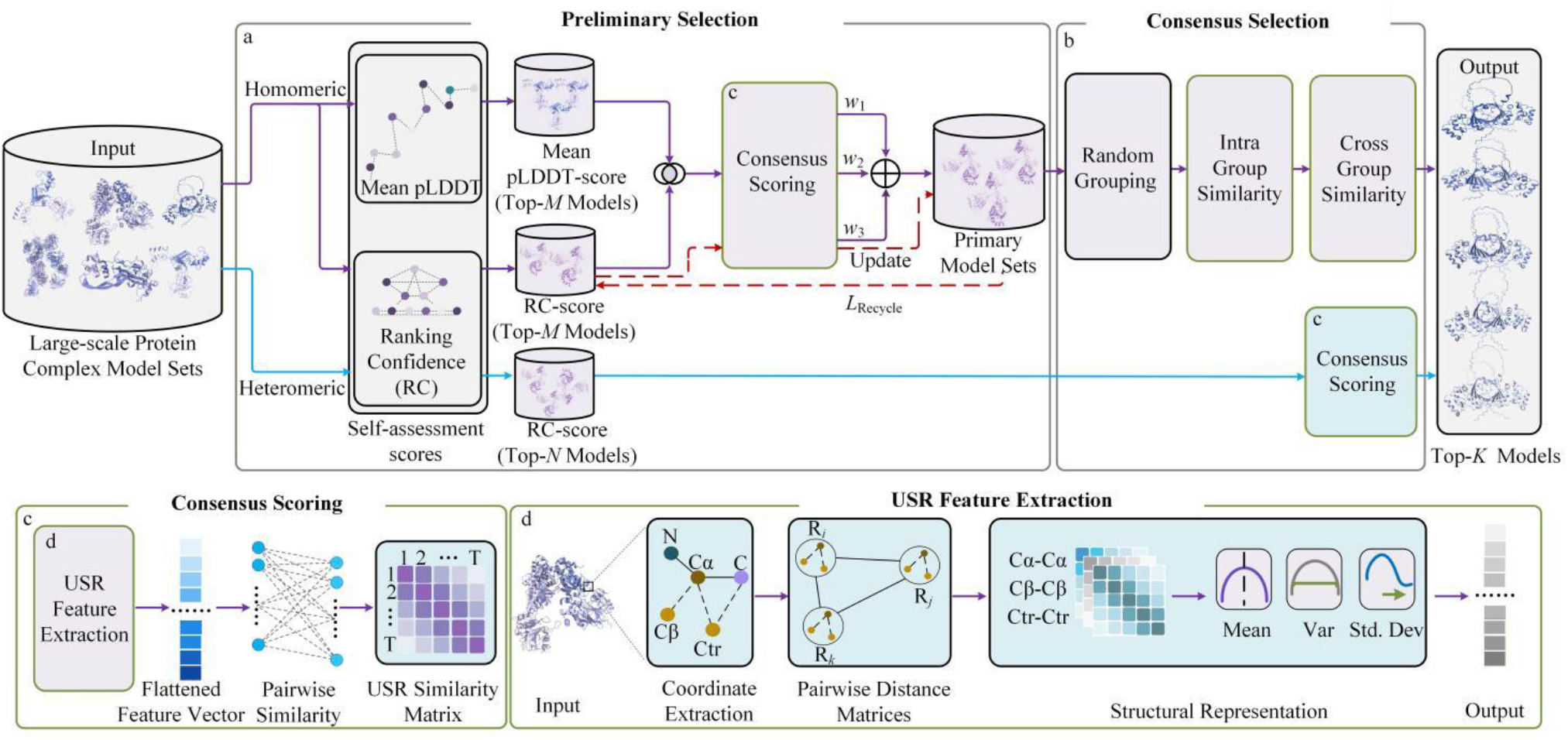
Overview of the confidence-assisted DeepUMQA-Selection workflow for large protein-complex targets. For large protein-complex targets with sequence lengths >1,500 residues, predictor-derived confidence metrics are incorporated into the preliminary selection stage by replacing the model-agnostic single-model quality estimators used in the standard DeepUMQA-Selection workflow, whereas the USR-based structural consensus and downstream selection procedures remain unchanged. The final ranked Top-*K* models are returned as the selected candidates. **(a) Preliminary selection**. For large homomeric complexes, Mean pLDDT and AlphaFold-Multimer ranking confidence (0.8 × ipTM + 0.2 × pTM) are used as two confidence scores to rank candidate models, replacing EMA_SGT_ and EMA_SIT_, respectively. The Top-*M* models from each ranking are retained, and their intersection is subjected to USR-based consensus scoring. The normalized mean pLDDT, ranking confidence, and USR consensus scores are combined using weights *w*_1_, *w*_2_, and *w*_3_, respectively, to construct the primary model set, which is subsequently refined through iterative replacement. For large heteromeric complexes, AlphaFold-Multimer ranking confidence replaces EMA_SIT_ for preliminary selection. The Top-*N* models identified by ranking confidence are used as references for subsequent consensus ranking. **(b) Consensus selection**. The branch-specific consensus-selection procedures are identical to those in the standard DeepUMQA-Selection workflow. For homomeric complexes, the refined primary model set is randomly partitioned into groups and ranked using hierarchical intra-group and inter-group USR consensus. For heteromeric complexes, candidate models are ranked according to their mean USR similarity to the reference models. **(c) Consensus scoring**. Structural similarity between two models is calculated from the Euclidean distance between their flattened USR feature vectors. A model’s consensus score is derived from its USR similarities to the models in the corresponding comparison set. **(d) USR feature extraction**. Residue-level C_α_, reconstructed C_β_, and centroid coordinates (Ctr) define three coordinate representations. For each representation, distance distributions associated with the residue itself, the residue farthest from it, and the residue farthest from that farthest residue are summarized by their mean (Mean), variance (Var), and standard deviation (Std.Dev), producing a nine-dimensional descriptor. The three descriptors are concatenated to generate a 27-dimensional USR vector for each residue.

### Supplementary Tables

**Supplementary Table S1.**
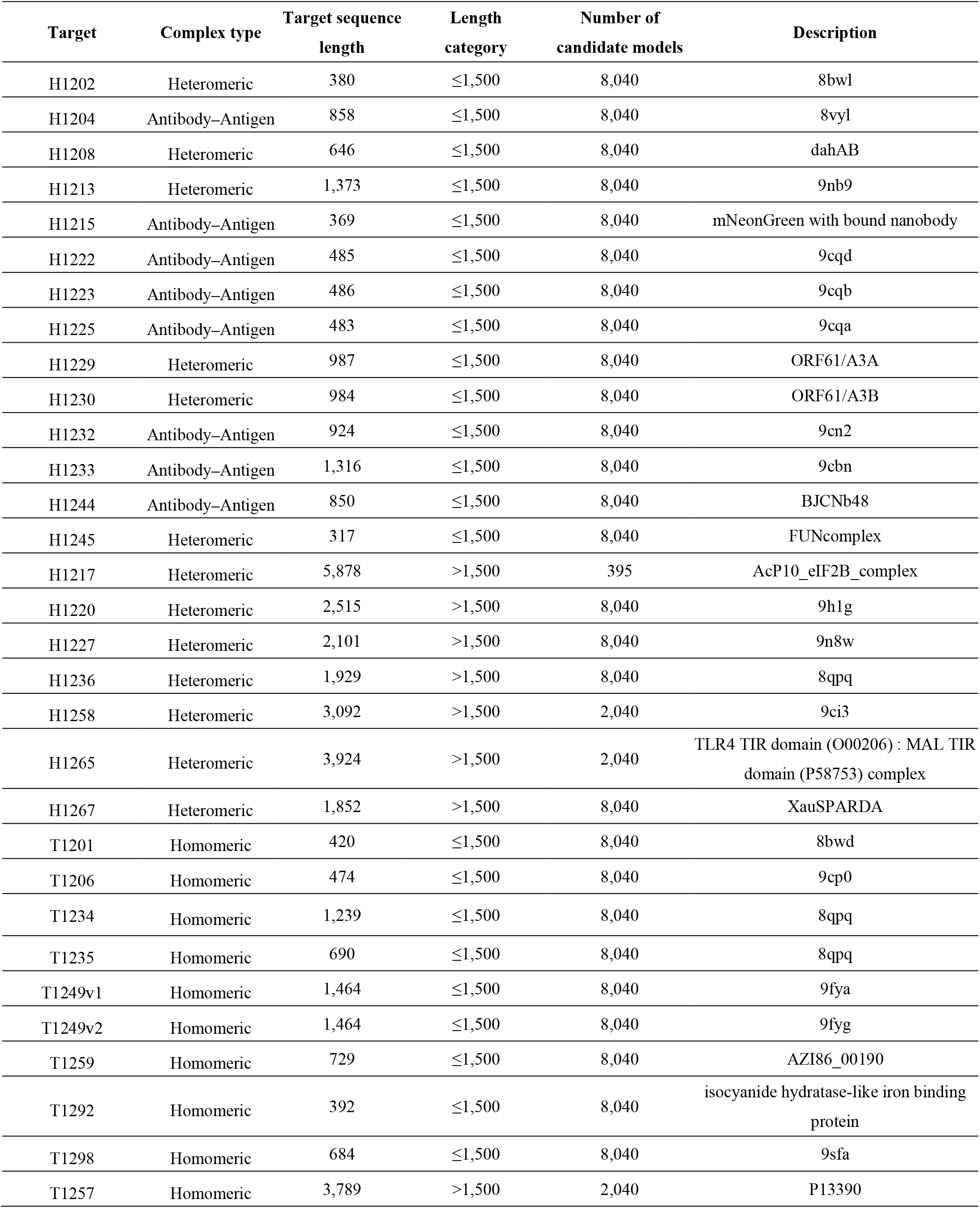
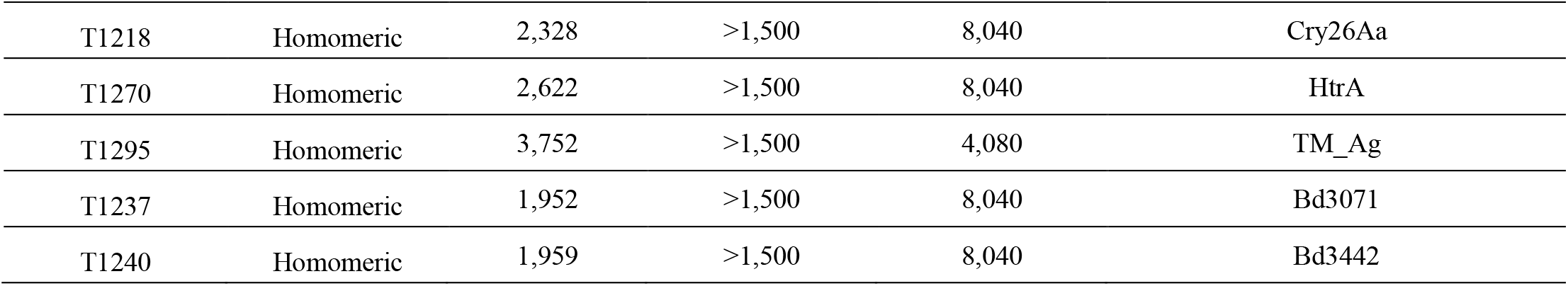
Summary of CASP16 QMODE3 protein complex targets and MassiveFold-generated candidate model pools. Target descriptions were obtained from the official CASP16 target list. Additional target information and associated metadata are available from the CASP16 Prediction Center website (https://predictioncenter.org/casp16/targetlist.cgi).

**Supplementary Table S2.**
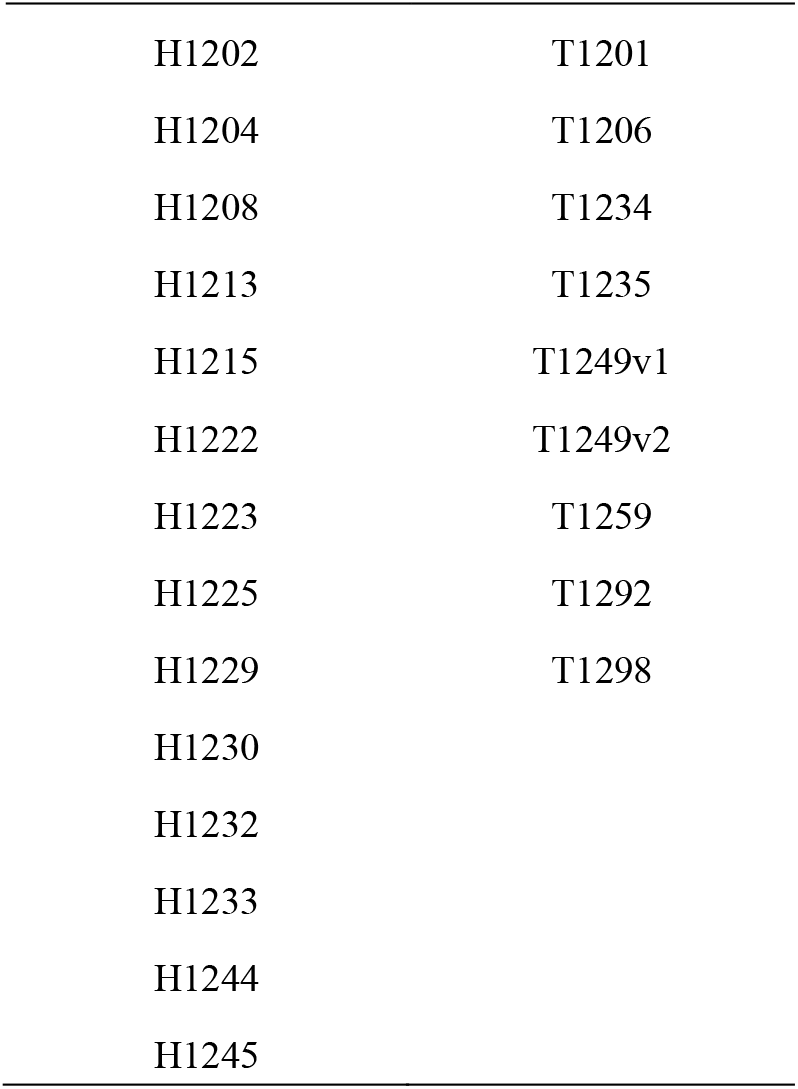
CASP16 QMODE3 targets with sequence lengths ≤1,500 residues.

**Supplementary Table S3.** CASP16 QMODE3 targets with sequence lengths >1,500 residues.

|  |  |
| --- | --- |
| H1217 | T1257 |
| H1220 | T1218 |
| H1227 | T1270 |
| H1236 | T1295 |
| H1258 | T1237 |
| H1265 | T1240 |
| H1267 |  |

**Supplementary Table S4.**
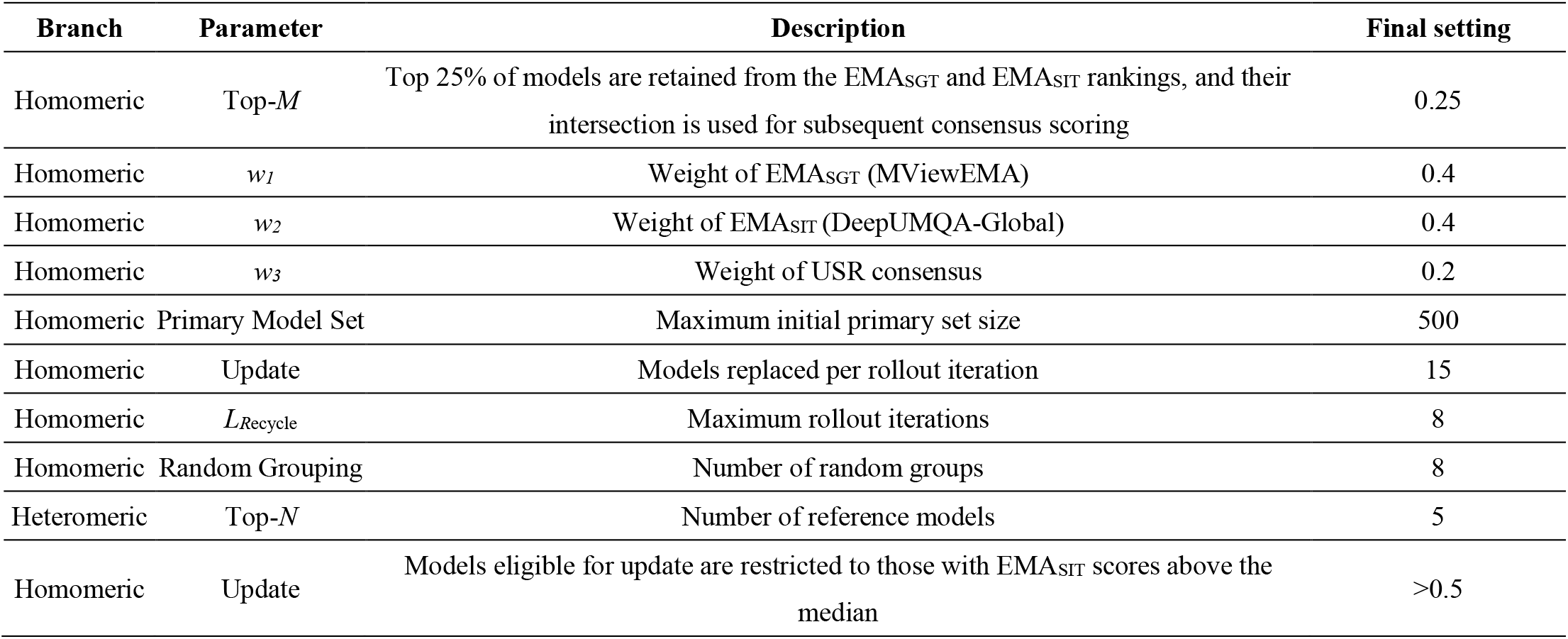
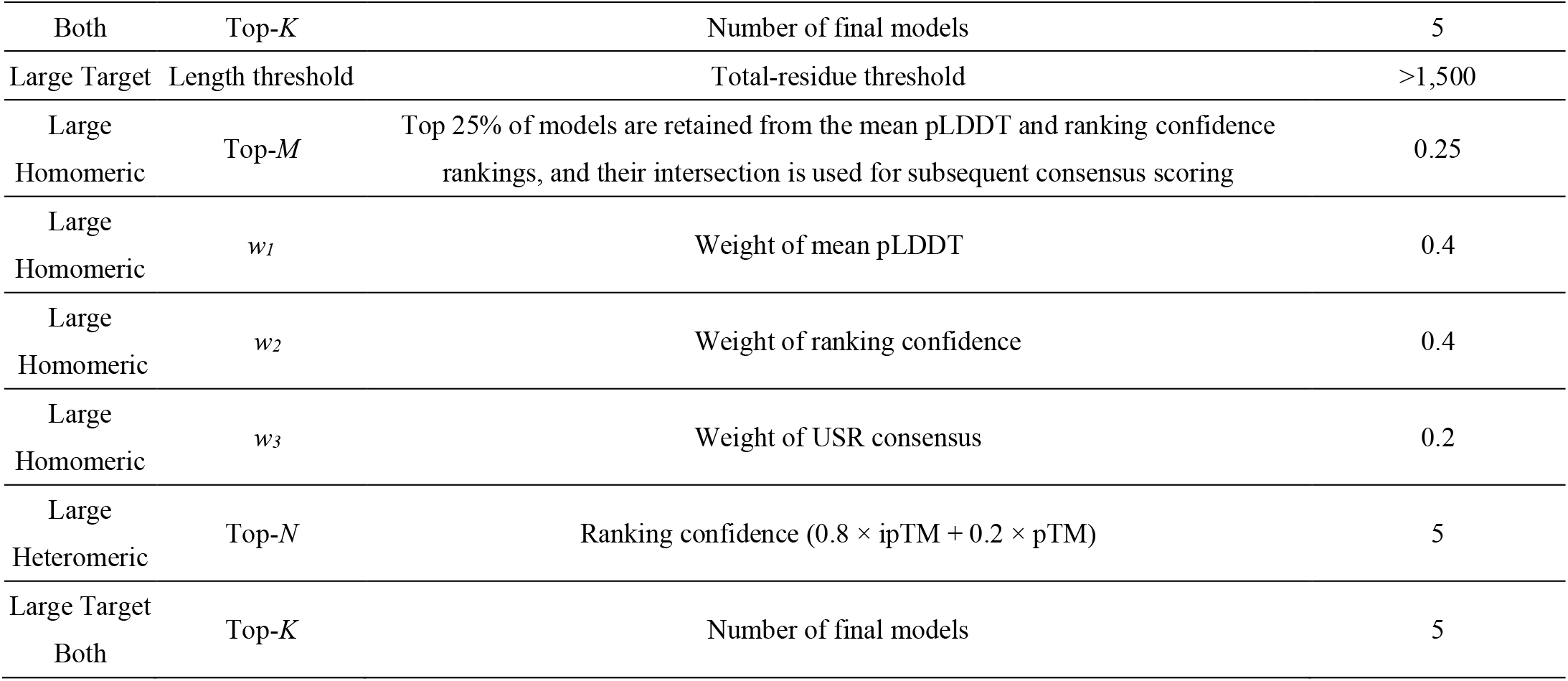
All parameter settings and branch-specific configurations of DeepUMQA-Selection.

| Branch | Parameter | Description | Final setting |
| --- | --- | --- | --- |
| Homomeric | Top-$M$ | Top 25% of models are retained from the EMA<sub>SGT</sub> and EMA<sub>SIT</sub> rankings, and their intersection is used for subsequent consensus scoring | 0.25 |
| Homomeric | $w_1$ | Weight of EMA<sub>SGT</sub> (MViewEMA) | 0.4 |
| Homomeric | $w_2$ | Weight of EMA<sub>SIT</sub> (DeepUMQA-Global) | 0.4 |
| Homomeric | $w_3$ | Weight of USR consensus | 0.2 |
| Homomeric | Primary Model Set | Maximum initial primary set size | 500 |
| Homomeric | Update | Models replaced per rollout iteration | 15 |
| Homomeric | $L_{Recycle}$ | Maximum rollout iterations | 8 |
| Homomeric | Random Grouping | Number of random groups | 8 |
| Heteromeric | Top-$N$ | Number of reference models | 5 |
| Homomeric | Update | Models eligible for update are restricted to those with EMA<sub>SIT</sub> scores above the median | $>0.5$ |
| Both | Top-$K$ | Number of final models | 5 |
| Large Target | Length threshold | Total-residue threshold | >1,500 |
| Large Homomeric | Top-$M$ | Top 25% of models are retained from the mean pLDDT and ranking confidence rankings, and their intersection is used for subsequent consensus scoring | 0.25 |
| Large Homomeric | $w_1$ | Weight of mean pLDDT | 0.4 |
| Large Homomeric | $w_2$ | Weight of ranking confidence | 0.4 |
| Large Homomeric | $w_3$ | Weight of USR consensus | 0.2 |
| Large Heteromeric | Top-$N$ | Ranking confidence ($0.8 \times \text{ipTM} + 0.2 \times \text{pTM}$) | 5 |
| Large Target Both | Top-$K$ | Number of final models | 5 |

**Supplementary Table S5.** Numerical data underlying the weighted penalty comparisons in Figure 2a–d.

(a) Group-level weighted-penalty comparisons (Figure 2a,b).
| Group (Heteromeric) | Weighted Penalty(↓) | Group (Homomeric) | Weighted Penalty(↓) |
| --- | --- | --- | --- |
| **Ours** | **0.229** | **Ours** | **0.113** |
| QA148 | 0.266 | QA331 | 0.114 |
| QA164 | 0.274 | QA050 | 0.127 |
| QA312 | 0.313 | QA051 | 0.130 |
| QA110 | 0.317 | QA345 | 0.145 |
| QA476 | 0.318 | QAptm | 0.149 |
| QA050 | 0.329 | QAiptm+ptm | 0.150 |
| QA264 | 0.337 | QAiptm | 0.150 |
| QA393 | 0.350 | QAmean_plddt | 0.156 |
| QA188 | 0.351 | QA314 | 0.166 |
| QAptm | 0.359 | QA319 | 0.185 |
| QA443 | 0.363 | QA273 | 0.213 |
| QAmean_plddt | 0.372 | QA122 | 0.214 |
| QAiptm+ptm | 0.381 | QA443 | 0.227 |
| QAiptm | 0.382 | QA148 | 0.227 |
| QA425 | 0.389 | QA425 | 0.234 |
| QA345 | 0.391 | QA029 | 0.248 |
| QA051 | 0.391 | QA110 | 0.310 |
| QA331 | 0.414 | QA393 | 0.321 |
| QA273 | 0.457 | QA312 | 0.334 |
| QA319 | 0.461 | QA264 | 0.370 |
| QA122 | 0.462 | QA212 | 0.393 |
| QA314 | 0.469 | QA188 | 0.490 |
| QA029 | 0.548 | QA476 | 0.537 |
| QA212 | 0.730 | QA164 | 0.609 |

| Target (Heteromeric) | Ours | QAiptm+ptm | Target (Homomeric) | Ours | QAiptm+ptm |
| --- | --- | --- | --- | --- | --- |
| H1202 | **0.030** | 0.100 | T1201 | **0.118** | 0.151 |
| H1204 | **0.109** | 0.998 | T1206 | **0.007** | 0.017 |
| H1208 | **0.021** | 0.583 | T1234 | **0.056** | 0.189 |
| H1213 | **0.005** | 0.085 | T1235 | 0.446 | **0.000** |
| H1215 | 0.730 | **0.194** | T1249v1 | **0.015** | 0.029 |
| H1222 | 0.718 | **0.440** | T1249v2 | **0.001** | 0.023 |
| H1223 | **0.016** | 0.594 | T1259 | **0.171** | 0.305 |
| H1225 | **0.000** | 0.283 | T1292 | **0.203** | 0.209 |
| H1229 | 0.743 | **0.503** | T1298 | **0.000** | 0.424 |
| H1230 | **0.124** | 0.321 |  |  |  |
| H1232 | **0.289** | 0.354 |  |  |  |
| H1233 | 0.289 | **0.150** |  |  |  |
| H1244 | **0.052** | 0.095 |  |  |  |
| H1245 | **0.074** | 0.630 |  |  |  |

**Supplementary Table S6.**
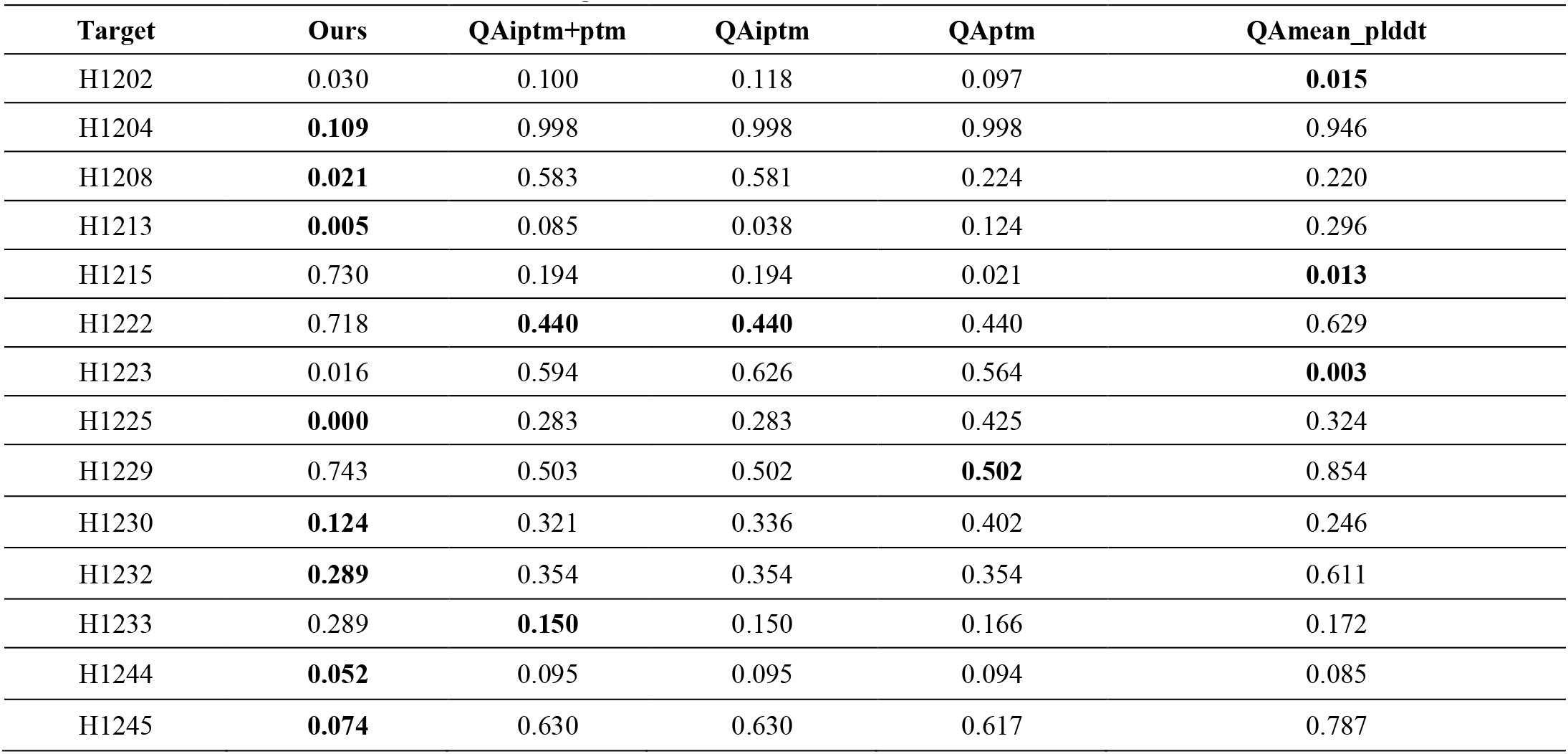
Target-wise weighted penalty comparison of DeepUMQA-Selection and AlphaFold-Multimer -assessment on CASP16 heteromeric targets.

**Supplementary Table S7.**
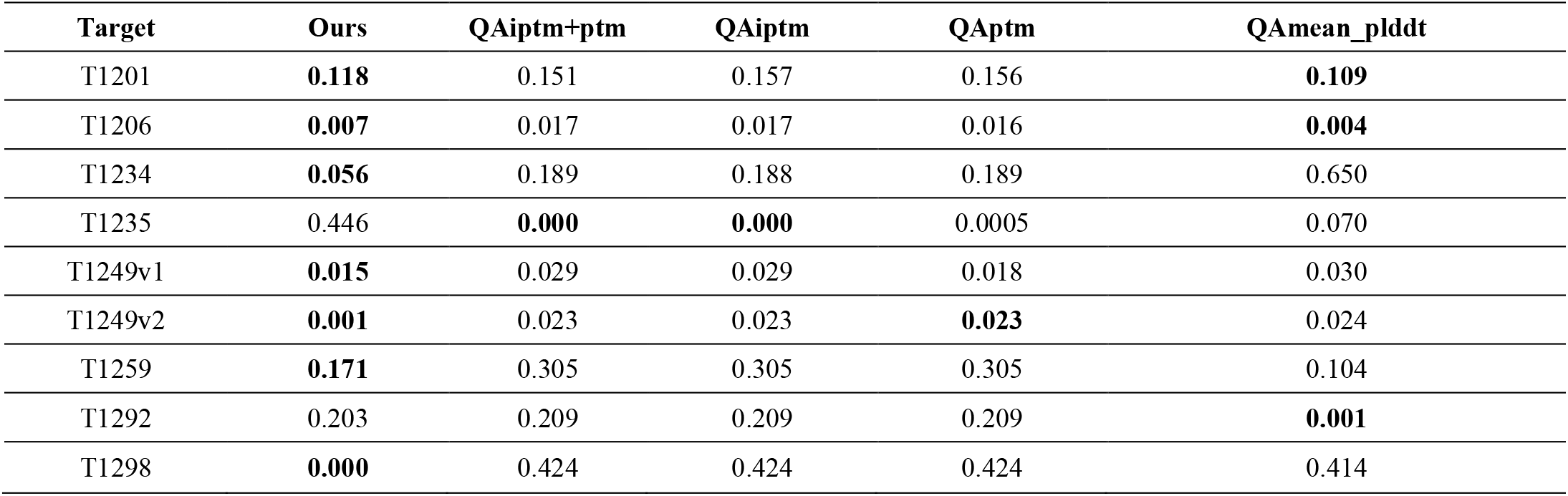
Target-wise weighted penalty comparison of DeepUMQA-Selection and AlphaFold-Multimer self-assessment on CASP16 homomeric targets.

**Supplementary Table S8.**
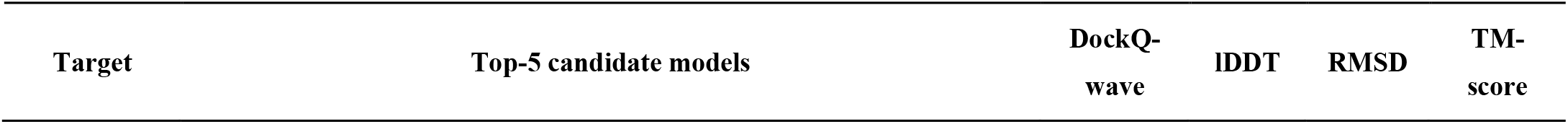
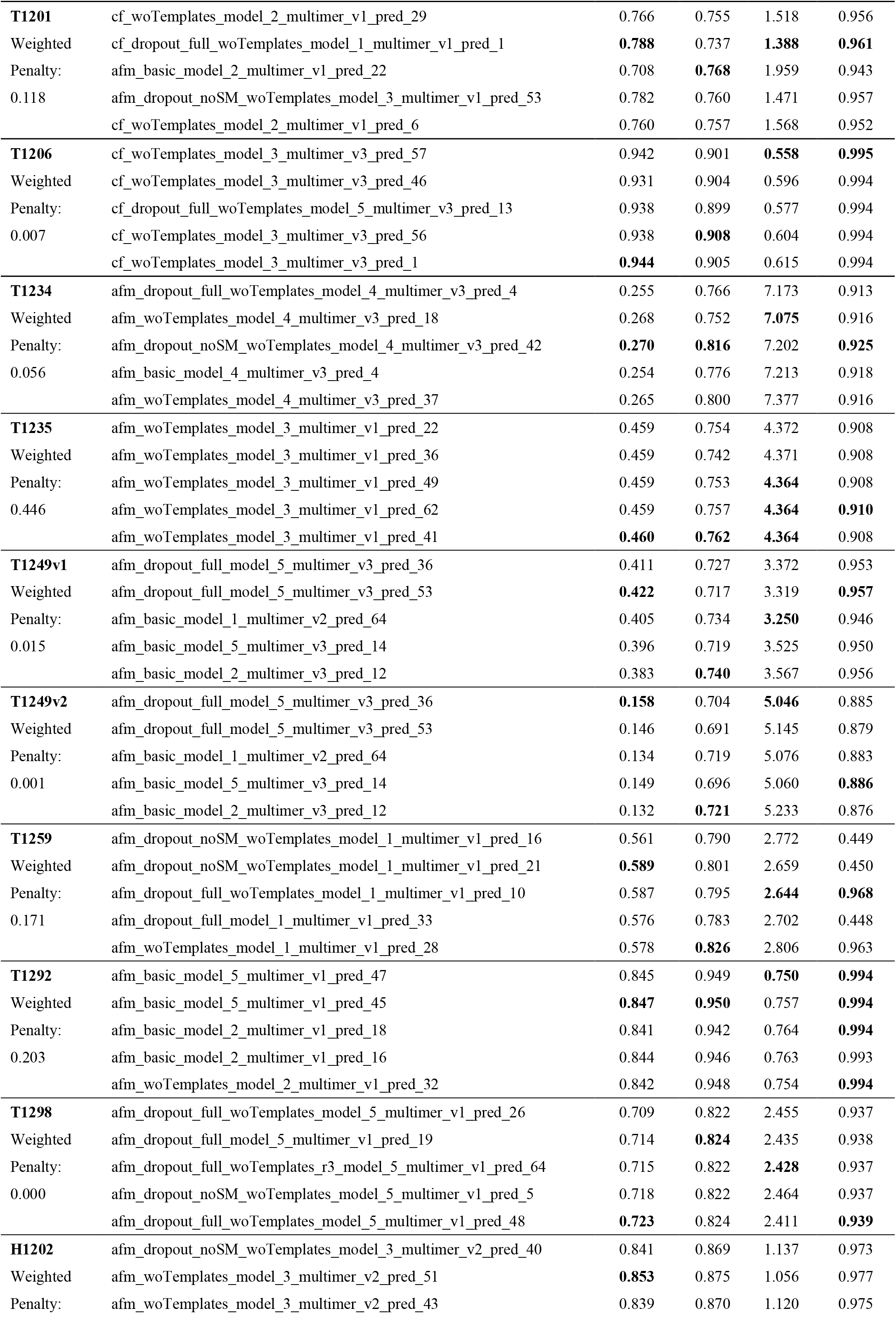
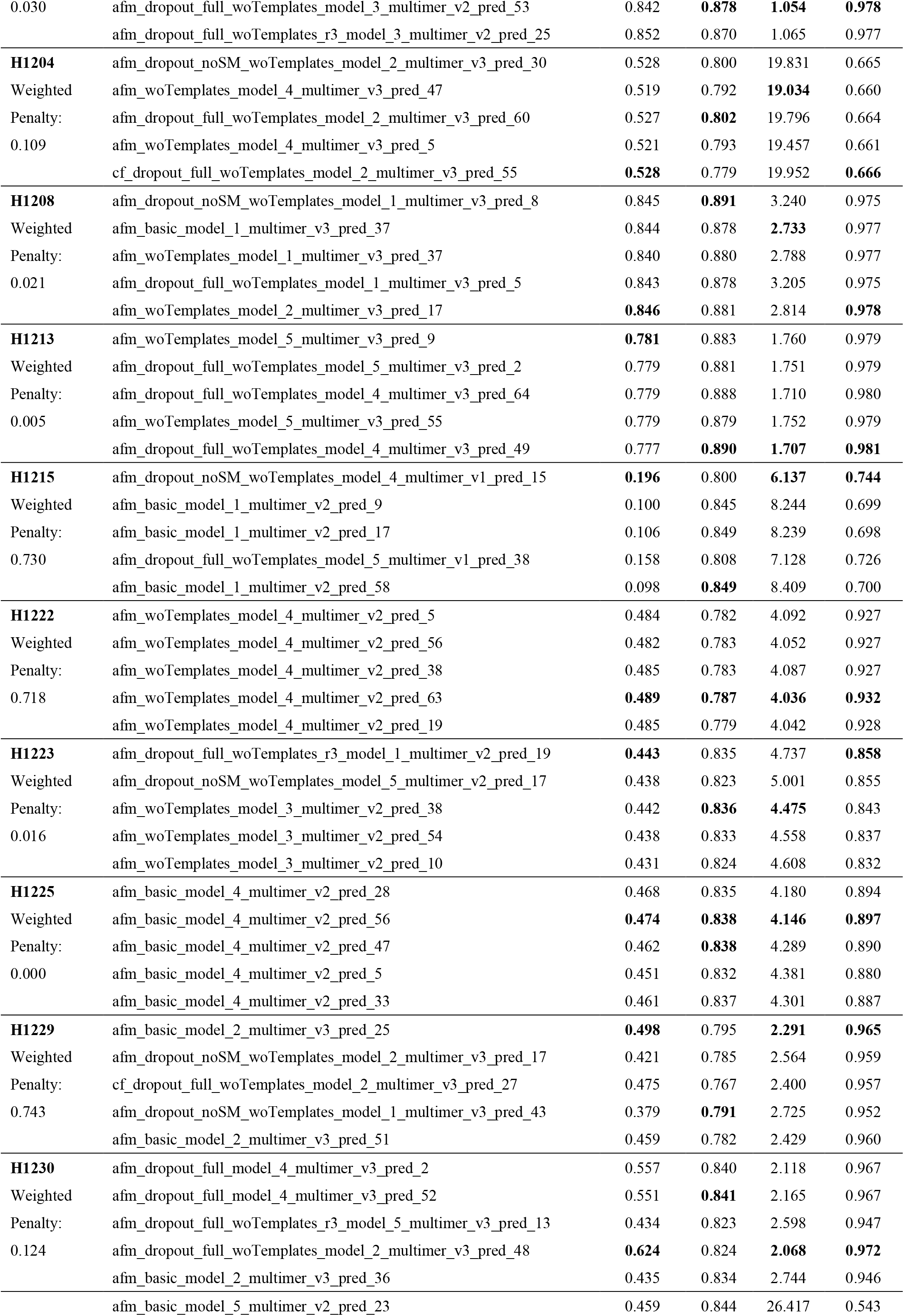
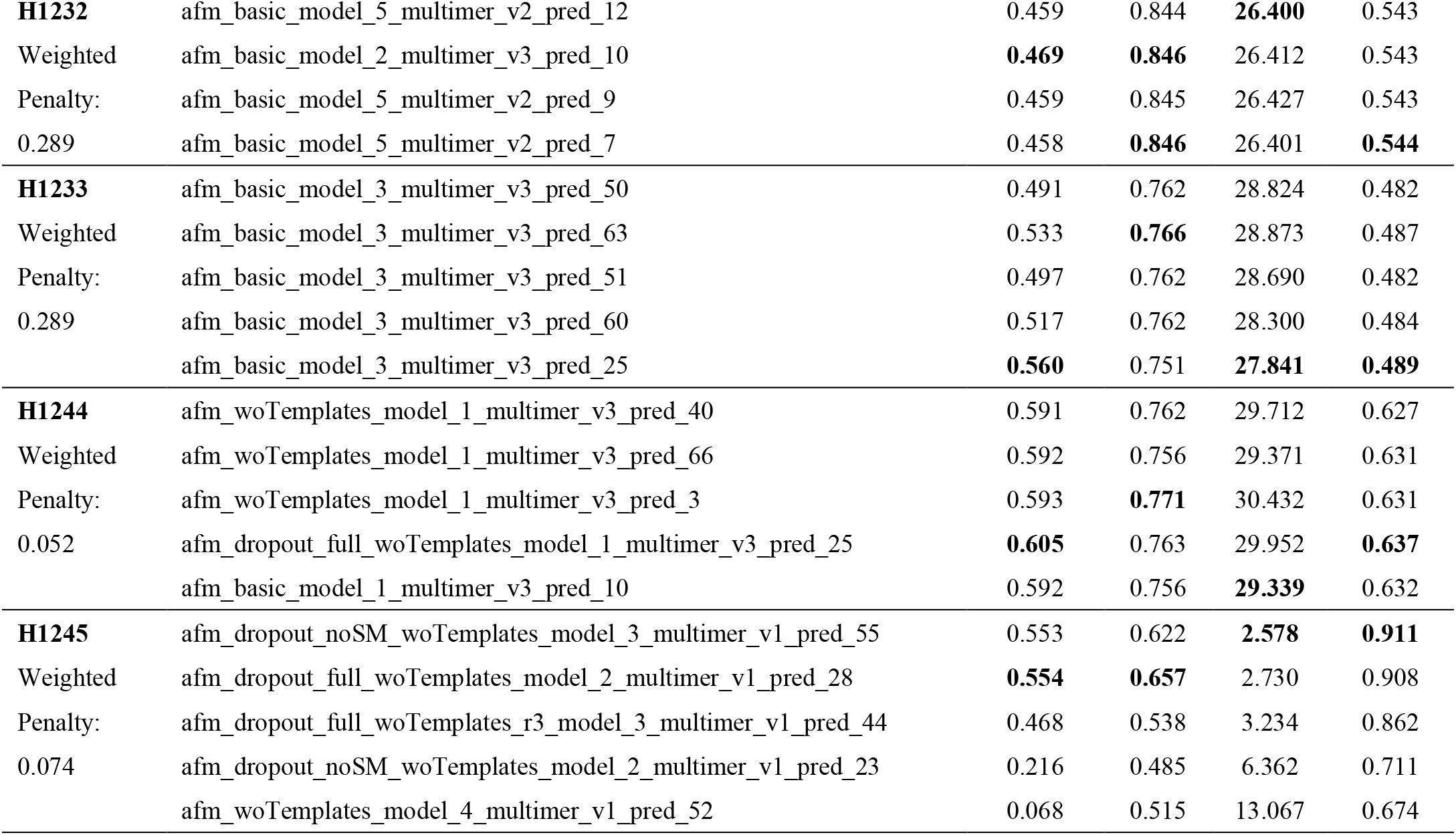
Top-5 candidate models selected by DeepUMQA-Selection and their weighted penalties and evaluation metric values for CASP16 homomeric and heteromeric targets with sequence lengths ≤1,500 residues.

**Supplementary Table S9.** Top-5 candidate models selected by DeepUMQA-Selection and their weighted penalties and evaluation metric values for CASP16 homomeric and heteromeric targets with sequence lengths >1,500 residues.

| Target | Top-5 candidate models | DockQ-wave | IDDT | RMSD | TM-score |
| --- | --- | --- | --- | --- | --- |
| **T1218** | afm_dropout_full_woTemplates_model_2_multimer_v3_pred_49 | **0.002** | 0.759 | 45.814 | 0.322 |
| Weighted | afm_dropout_noSM_woTemplates_model_2_multimer_v3_pred_18 | **0.002** | 0.760 | **45.556** | 0.318 |
| Penalty: | afm_dropout_full_woTemplates_model_2_multimer_v3_pred_31 | **0.002** | 0.762 | 45.666 | **0.326** |
| 0.023 | afm_dropout_noSM_woTemplates_model_2_multimer_v3_pred_16 | **0.002** | **0.764** | 45.846 | 0.310 |
|  | afm_dropout_full_woTemplates_model_2_multimer_v3_pred_24 | **0.002** | 0.762 | 45.826 | 0.317 |
| **T1237** | afm_woTemplates_model_1_multimer_v3_pred_45 | 0.775 | 0.865 | **2.227** | **0.976** |
| Weighted | afm_woTemplates_model_1_multimer_v3_pred_57 | 0.774 | 0.873 | 2.277 | 0.975 |
| Penalty: | afm_woTemplates_model_1_multimer_v3_pred_60 | 0.774 | **0.879** | 2.270 | 0.975 |
| 0.034 | afm_woTemplates_model_1_multimer_v3_pred_25 | 0.774 | 0.872 | 2.237 | 0.975 |
|  | afm_basic_model_1_multimer_v3_pred_59 | **0.776** | 0.878 | 2.298 | 0.974 |
| **T1240** | afm_dropout_full_woTemplates_model_1_multimer_v3_pred_61 | 0.316 | 0.806 | 10.704 | 0.631 |
| Weighted | afm_dropout_noSM_woTemplates_model_1_multimer_v3_pred_15 | **0.322** | 0.806 | **10.388** | **0.643** |
| Penalty: | afm_dropout_noSM_woTemplates_model_1_multimer_v3_pred_54 | 0.319 | 0.801 | 10.600 | 0.638 |
| 0.526 | afm_dropout_noSM_woTemplates_model_1_multimer_v3_pred_55 | 0.317 | **0.810** | 10.531 | 0.643 |
|  | afm_dropout_noSM_woTemplates_model_1_multimer_v3_pred_9 | 0.307 | 0.804 | 10.740 | 0.629 |
| **T1257** | afm_woTemplates_model_5_multimer_v3_pred_8 | 0.253 | 0.685 | 117.947 | **0.664** |
| Weighted | cf_woTemplates_model_5_multimer_v3_pred_14 | 0.252 | 0.705 | 116.323 | 0.537 |
| Penalty: | afm_woTemplates_model_5_multimer_v3_pred_3 | **0.263** | **0.724** | 98.409 | 0.578 |
| 0.138 | afm_woTemplates_model_5_multimer_v3_pred_11 | 0.251 | 0.671 | 113.198 | 0.654 |
|  | afm_dropout_noSM_woTemplates_model_2_multimer_v3_pred_8 | 0.241 | 0.594 | **97.340** | 0.581 |
| **T1270** | afm_basic_model_4_multimer_v3_pred_33 | 0.576 | **0.801** | 17.489 | 0.506 |
| Weighted | afm_woTemplates_model_4_multimer_v3_pred_12 | 0.577 | 0.796 | 17.807 | 0.501 |
| Penalty: | afm_woTemplates_model_4_multimer_v3_pred_54 | 0.576 | 0.798 | 17.799 | 0.499 |
| 0.045 | afm_dropout_noSM_woTemplates_model_4_multimer_v3_pred_49 | 0.583 | 0.794 | **17.349** | 0.505 |
|  | afm_dropout_full_woTemplates_model_4_multimer_v3_pred_44 | **0.585** | 0.797 | 17.353 | **0.509** |
| **T1295** | cf_dropout_full_woTemplates_model_2_multimer_v3_pred_19 | 0.247 | 0.752 | **43.421** | 0.195 |
| Weighted | afm_dropout_noSM_woTemplates_model_2_multimer_v3_pred_33 | 0.242 | 0.756 | 52.512 | 0.164 |
| Penalty: | cf_dropout_full_woTemplates_model_2_multimer_v3_pred_3 | 0.252 | 0.765 | 52.727 | **0.201** |
| 0.015 | cf_dropout_full_woTemplates_model_2_multimer_v3_pred_20 | 0.242 | **0.766** | 52.063 | 0.193 |
|  | cf_dropout_full_woTemplates_model_2_multimer_v3_pred_32 | **0.255** | 0.754 | 52.751 | 0.200 |
| **H1217** | afm_dropout_full_model_5_multimer_v3_pred_0 | 0.675 | 0.757 | 29.605 | 0.855 |
| Weighted | afm_dropout_full_model_1_multimer_v3_pred_0 | **0.690** | 0.784 | **28.704** | **0.885** |
| Penalty: | afm_basic_model_5_multimer_v3_pred_4 | 0.682 | 0.755 | 32.810 | 0.854 |
| 0.848 | afm_basic_model_4_multimer_v3_pred_3 | 0.674 | **0.808** | 31.388 | 0.869 |
|  | afm_dropout_full_model_1_multimer_v3_pred_4 | 0.687 | 0.794 | 29.083 | 0.884 |
| **H1220** | cf_woTemplates_model_5_multimer_v3_pred_41 | 0.744 | 0.723 | 3.559 | **0.945** |
| Weighted | cf_woTemplates_model_5_multimer_v3_pred_10 | 0.741 | 0.726 | **3.540** | **0.945** |
| Penalty: | cf_woTemplates_model_5_multimer_v3_pred_44 | 0.742 | 0.725 | 3.788 | 0.939 |
| 0.008 | cf_woTemplates_model_5_multimer_v3_pred_22 | **0.754** | 0.725 | 3.587 | 0.943 |
|  | cf_woTemplates_model_5_multimer_v3_pred_8 | 0.753 | **0.730** | 3.691 | 0.941 |
| **H1227** | afm_basic_model_5_multimer_v3_pred_43 | 0.500 | 0.784 | 3.714 | 0.939 |
| Weighted | afm_basic_model_1_multimer_v3_pred_27 | 0.506 | 0.788 | 4.239 | 0.920 |
| Penalty: | afm_dropout_noSM_woTemplates_model_3_multimer_v3_pred_7 | **0.648** | **0.805** | **2.044** | **0.979** |
| 0.125 | afm_woTemplates_model_3_multimer_v3_pred_65 | 0.635 | 0.803 | 2.120 | 0.978 |
|  | afm_dropout_full_model_1_multimer_v3_pred_7 | 0.511 | 0.783 | 4.276 | 0.919 |
| **H1236** | afm_dropout_noSM_woTemplates_model_4_multimer_v3_pred_45 | **0.304** | **0.788** | **15.999** | **0.762** |
| Weighted | afm_dropout_noSM_woTemplates_model_4_multimer_v3_pred_22 | 0.291 | 0.786 | 16.134 | 0.756 |
| Penalty: | afm_dropout_full_model_5_multimer_v3_pred_30 | 0.295 | 0.779 | 16.013 | 0.761 |
| 0.015 | afm_woTemplates_model_5_multimer_v3_pred_35 | 0.289 | 0.775 | 16.081 | 0.759 |
|  | afm_dropout_full_model_5_multimer_v3_pred_23 | 0.285 | 0.783 | 16.068 | 0.759 |
| **H1258** | afm_basic_model_1_multimer_v3_pred_12 | 0.255 | 0.746 | 15.848 | 0.784 |
| Weighted | afm_basic_model_1_multimer_v3_pred_13 | 0.357 | 0.755 | 13.086 | 0.776 |
| Penalty: | afm_basic_model_1_multimer_v3_pred_16 | 0.175 | 0.736 | 11.874 | 0.795 |
| 0.225 | afm_basic_model_1_multimer_v3_pred_1 | 0.247 | 0.749 | **11.044** | **0.808** |
|  | afm_basic_model_1_multimer_v3_pred_11 | **0.373** | **0.758** | 12.444 | 0.799 |
| **H1265** | cf_dropout_full_woTemplates_model_5_multimer_v3_pred_7 | 0.197 | **0.654** | 40.535 | 0.279 |
| Weighted | afm_basic_model_1_multimer_v3_pred_14 | 0.088 | 0.540 | **39.733** | **0.332** |
| Penalty: | cf_dropout_full_woTemplates_model_1_multimer_v3_pred_13 | **0.200** | 0.591 | 45.839 | 0.246 |
| 0.071 | afm_dropout_noSM_woTemplates_model_5_multimer_v3_pred_10 | 0.178 | 0.544 | 42.234 | 0.273 |
|  | afm_dropout_noSM_woTemplates_model_1_multimer_v3_pred_4 | 0.170 | 0.443 | 43.474 | 0.284 |
| **H1267** | cf_woTemplates_model_5_multimer_v3_pred_63 | 0.224 | 0.203 | 14.393 | 0.896 |
| Weighted | cf_dropout_full_woTemplates_model_1_multimer_v3_pred_25 | 0.246 | 0.200 | 14.349 | 0.900 |
| Penalty: | afm_basic_model_4_multimer_v3_pred_36 | 0.185 | 0.198 | 14.356 | 0.872 |
| 0.004 | cf_woTemplates_model_1_multimer_v3_pred_45 | **0.248** | 0.204 | 14.371 | **0.902** |
|  | afm_dropout_noSM_woTemplates_model_1_multimer_v3_pred_4 | 0.231 | **0.207** | **14.275** | 0.899 |

**Supplementary Table S10.**
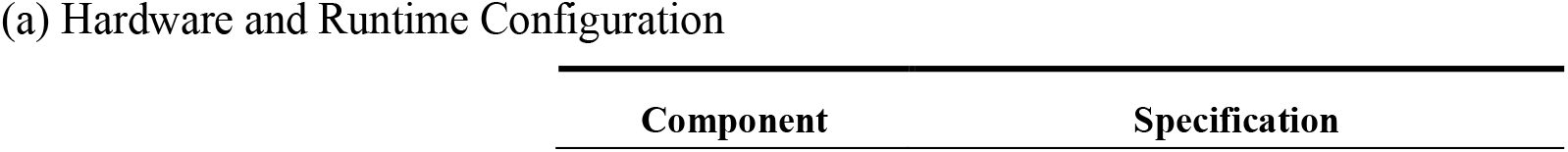
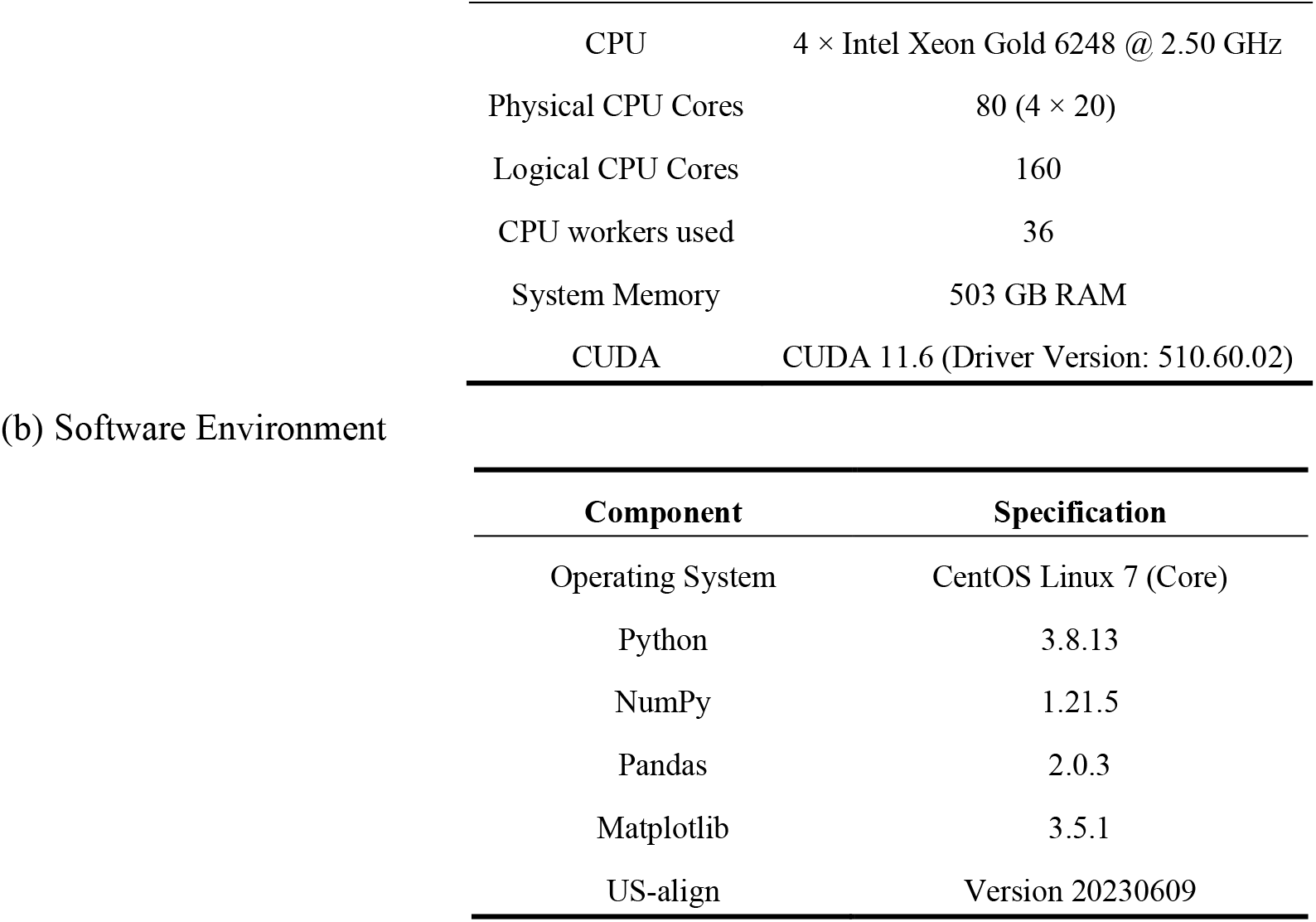
Hardware and software configuration of the compute node used for the efficiency experiments. The efficiency experiments were conducted on a single compute node of an in-house computing cluster. Only the hardware resources and software environment actually used in the benchmark are reported.

| Component | Specification |
| --- | --- |
| CPU | 4 × Intel Xeon Gold 6248 @ 2.50 GHz |
| Physical CPU Cores | 80 (4 × 20) |
| Logical CPU Cores | 160 |
| CPU workers used | 36 |
| System Memory | 503 GB RAM |
| CUDA | CUDA 11.6 (Driver Version: 510.60.02) |

(b) Software Environment
| Component | Specification |
| --- | --- |
| Operating System | CentOS Linux 7 (Core) |
| Python | 3.8.13 |
| NumPy | 1.21.5 |
| Pandas | 2.0.3 |
| Matplotlib | 3.5.1 |
| US-align | Version 20230609 |

### Supplementary Algorithms

#### Supplementary Algorithm 1.

Ultrafast Shape Recognition (USR)-Based Structural Alignment

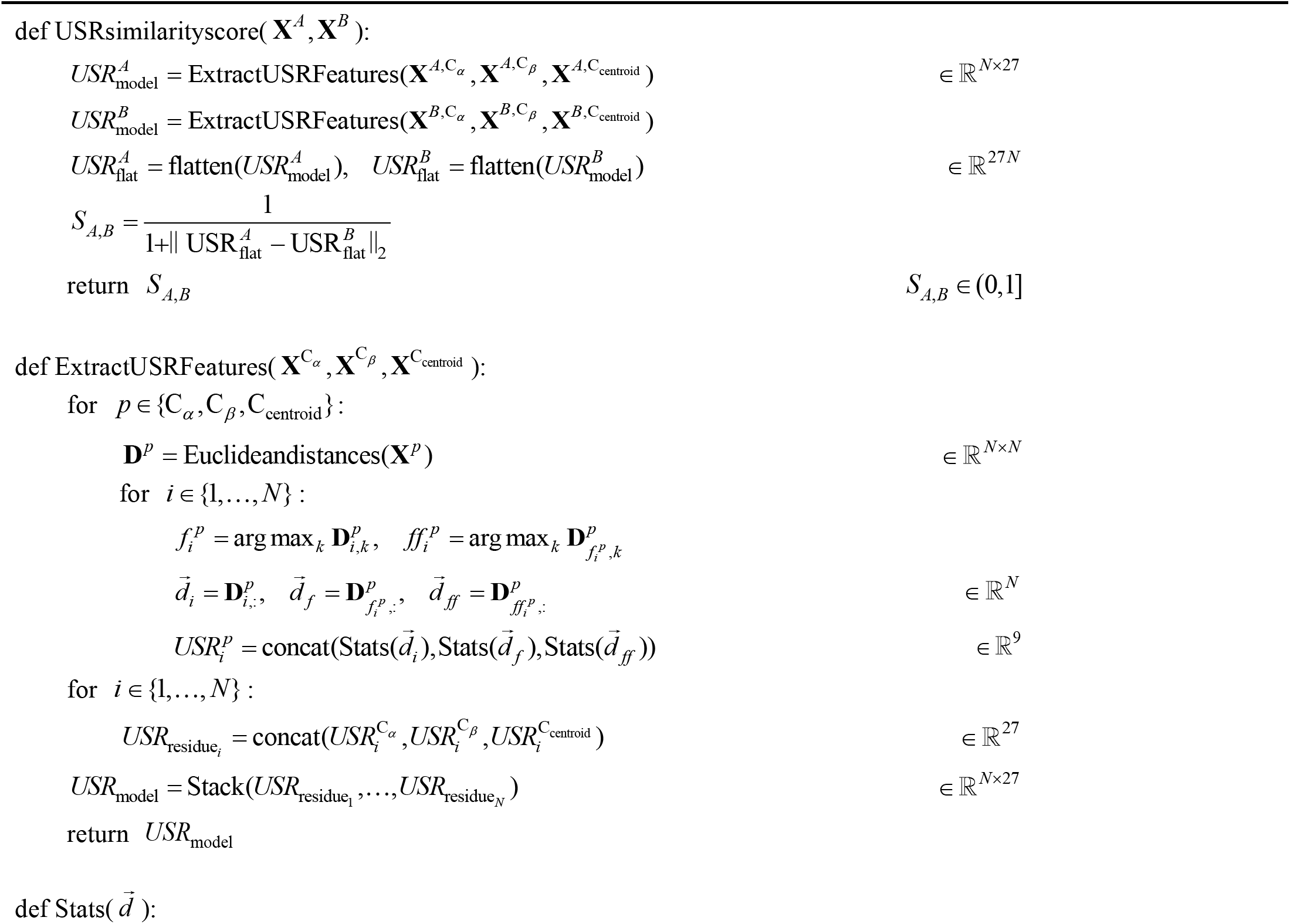
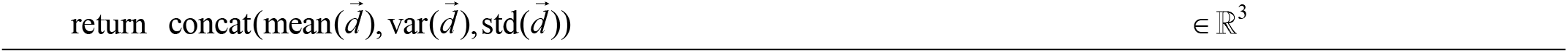

#### Supplementary Algorithm 2.

Primary model set mini-rollout update

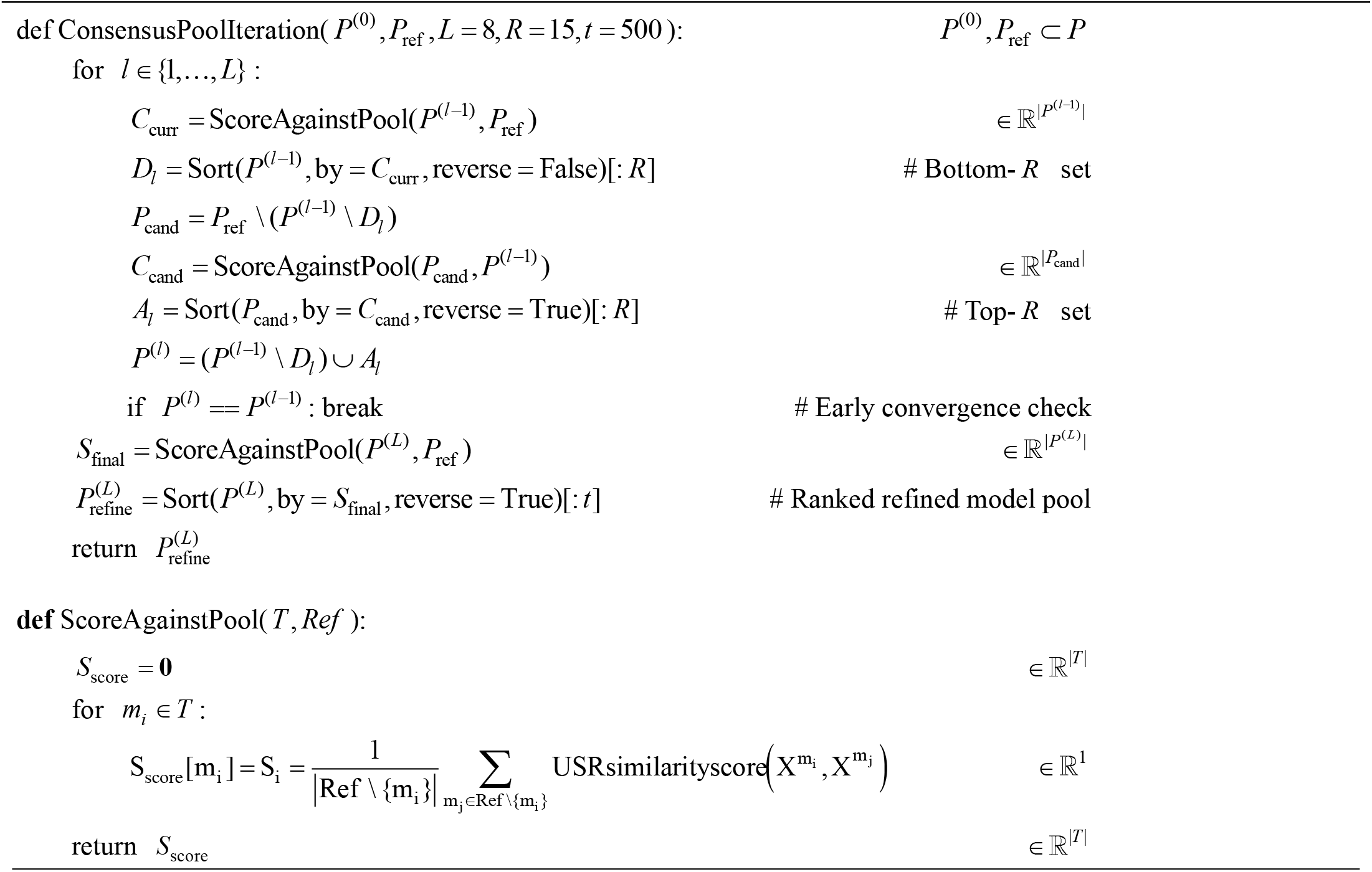

#### Supplementary Algorithm 3.

Hierarchical Consensus Selection

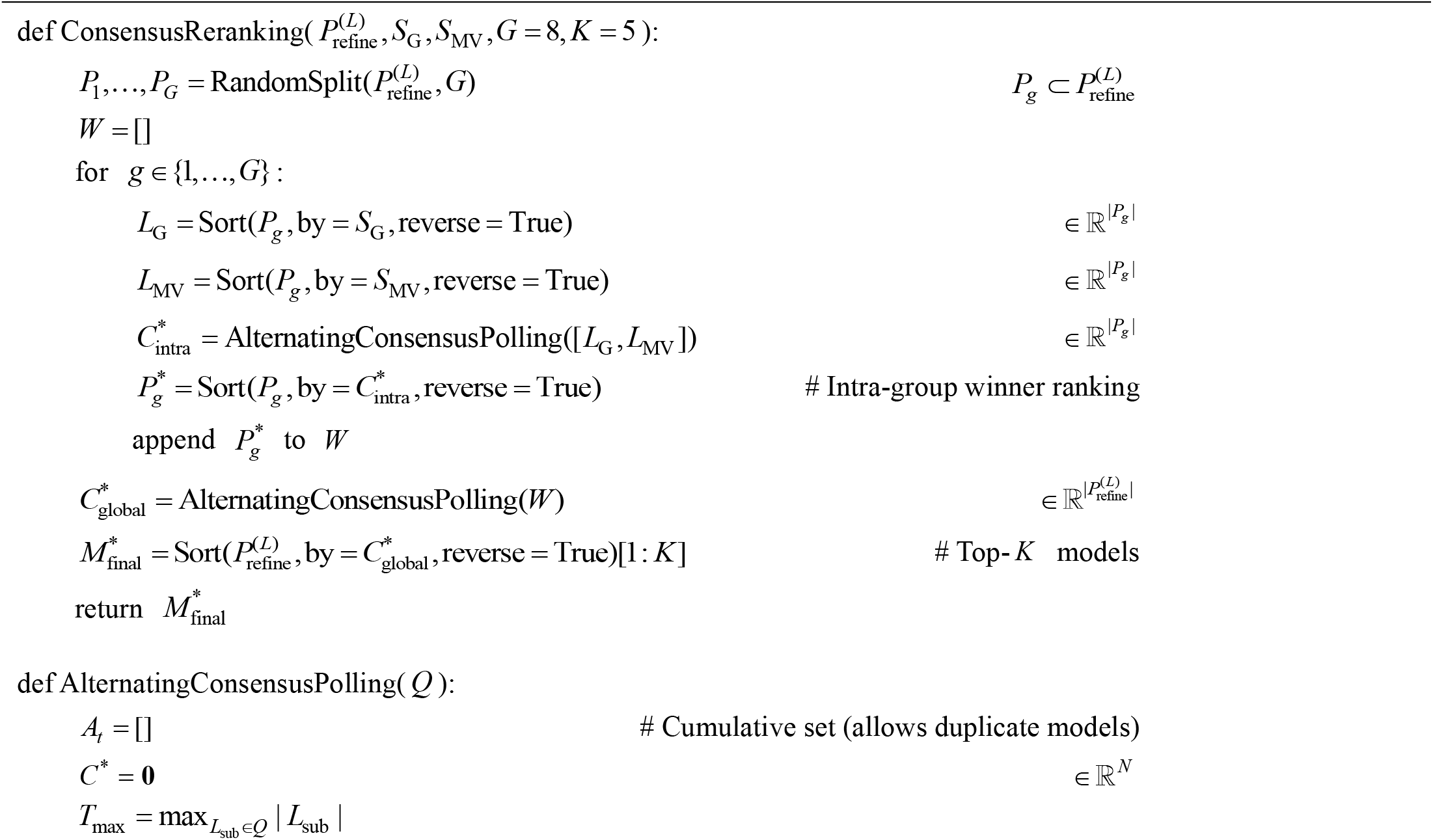
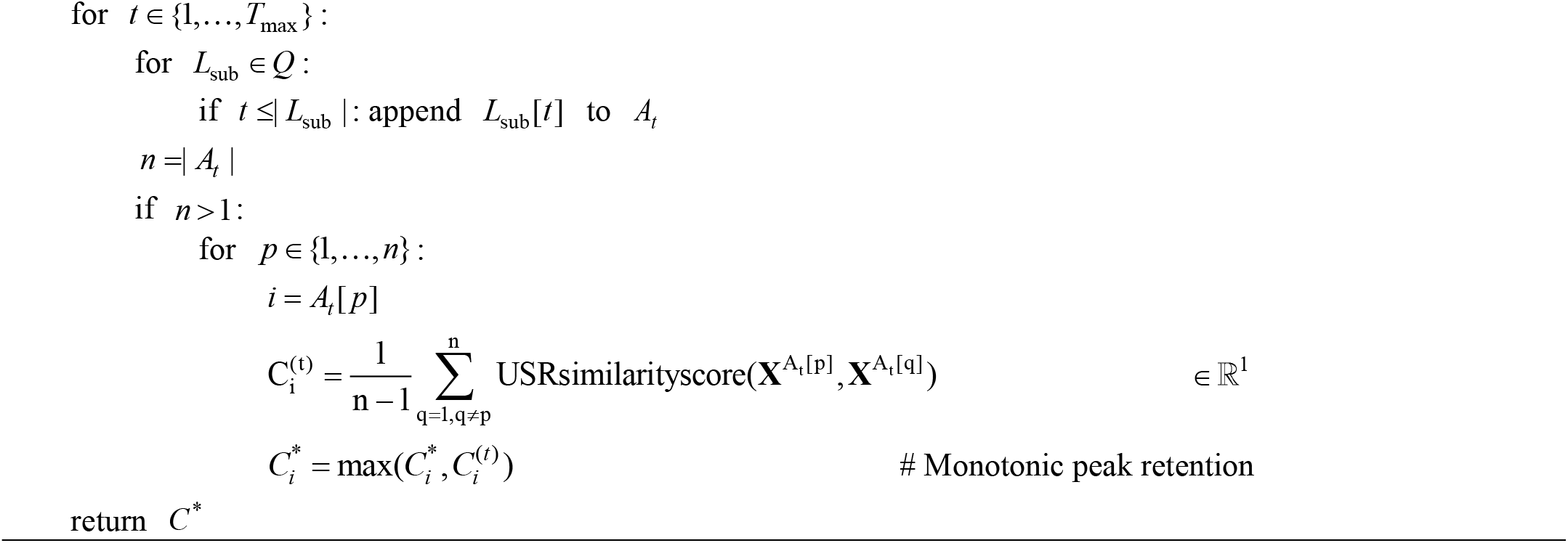

### Supplementary Notes

#### Supplementary Note 1

Definition of evaluation metrics

Following the CASP16 QMODE3 assessment, OpenStructure (OST) was used to compute structural quality scores by comparing each model with its corresponding experimental target. lDDT, TM-score, QS-global, QS-best, RMSD, oligo-GDTHA, and oligo-GDTTS were calculated for all targets, whereas iLDDT, IPS, ICS, DockQ-ave, and DockQ-wave were additionally calculated for oligomeric targets. RMSD, interface contact score (ICS) and interface patch score (IPS) were not included in the weighted-penalty calculation.

##### TM-score

TM-score evaluates the similarity in overall fold and global topology between a candidate model and the reference structure. It uses a structure-length-dependent normalization factor to reduce the influence of large local deviations and protein length on the overall assessment. The metric is defined as[1]:

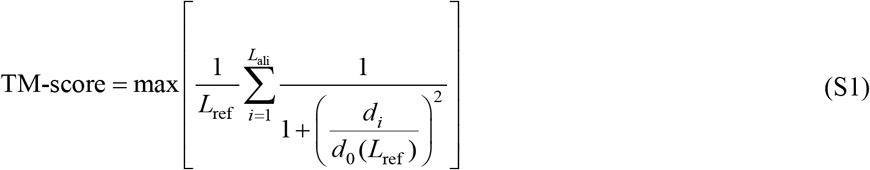

where *L*_ref_ is the length of the reference structure, *L*_ali_is the number of aligned residues after structural superposition, is the distance between the *i*-th pair of corresponding residues after superposition, and *d*_0_(*L*_ref_) is the normalization factor determined by the reference-structure length. TM-score generally ranges from 0 to 1, with higher values indicating greater similarity between the candidate model and the reference structure at the level of global topology.

##### lDDT

lDDT (Local Distance Difference Test) evaluates how well a predicted model preserves the local distance relationships between corresponding atom pairs in a reference structure. It does not require global structural superposition; instead, it calculates the fraction of local distances preserved within predefined tolerance thresholds and averages these fractions to obtain the final score. The metric is defined as[2,3]:

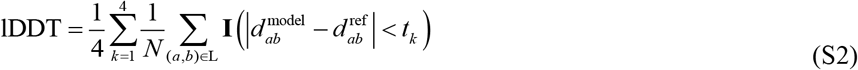

where L is the set of all atom pairs in the reference structure whose interatomic distance is less than the inclusion radius =15 Å and whose atoms belong to different residues; *N* = | L | is the total number of reference atom pairs included in the evaluation; 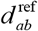 and 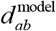 are the interatomic distances in the reference structure and the candidate model, respectively; *tk* denotes the four tolerance thresholds of 0.5, 1.0, 2.0, and 4.0I(Å·); and is the indicator function, which equals 1 when the corresponding distance difference is smaller than the specified threshold and 0 otherwise. If one or both atoms defining a reference distance are missing from the candidate model, that distance is considered non-preserved. The final lDDT score is the average fraction of preserved distances over the four tolerance thresholds and ranges from 0 to 1, with higher values indicating better preservation of local structural geometry.

##### iLDDT

iLDDT (The interface Local Distance Difference Test) is a superposition-free metric for evaluating the accuracy of protein–protein interfaces. As an interface-focused variant of lDDT, iLDDT measures how well local interatomic distance relationships across different chains in the reference structure are preserved in the candidate model. Only inter-chain atom pairs within an inclusion radius *R*_0_ =15 Å are considered. The metric is defined as[4]:

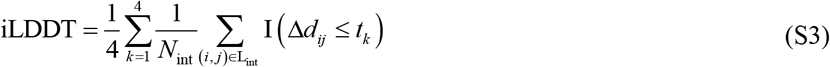

where Lint is the set of inter-chain atom pairs within the inclusion radius *R*_0_ =15 Å; *N*_int_ =∣L_in_∣_t_ is the total number of evaluated inter-chain atom pairs; 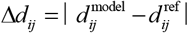 is the absolute difference between the corresponding interatomic distances in the candidate model and the reference structure; *t*_*k*_ denotes the four tolerance thresholds of 0.5, 1.0, 2.0, and Å; and I (·) is the indicator function, which equals 1 when the condition is satisfied and 0 otherwise. iLDDT ranges from 0 to 1, with higher values indicating better preservation of the interfacial geometry between interacting protein chains.

##### QS-score

QS-score evaluates the similarity between a predicted protein complex and a reference structure by considering both the conservation of inter-chain contacts and the corresponding inter-residue distance deviations. It provides 2 complementary variants, QS-global and QS-best, which can be expressed in a unified form as[5]:

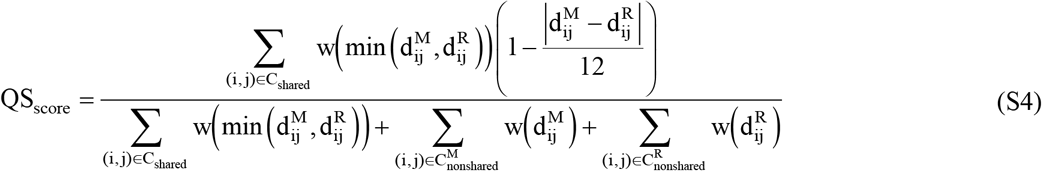

where 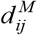 and 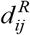 denote the distances between residues *ij* and in the candidate model and the reference structure, respectively; *C*_shared_ is the set of interchain contacts shared by the two structures; and 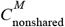 and 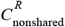 denote the corresponding sets of nonshared contacts. QS-global considers all nonshared contacts, whereas QS-best considers only nonshared contacts within successfully mapped chains and residues. Consequently, QS-best is generally greater than or equal to QS-global.

##### oligo-GDTTS

oligo-GDTTS is the oligomeric extension of the Global Distance Test–Total Score (GDT-TS)[6]. It evaluates the global coordinate accuracy of an entire complex under the optimal chain mapping and structural superposition.

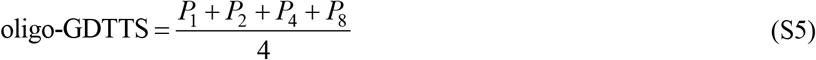

where *P*_1_, *P*_2_, *P*_4_ and *P*_8_ denote the fractions of aligned residue pairs whose corresponding Cα atoms are separated by no more than 1, 2, 4, and 8 Å, respectively, after structural superposition. A higher oligo-GDTTS value indicates greater overall structural accuracy of the candidate model across the entire complex.

##### oligo-GDTHA

oligo-GDTHA is the oligomeric extension of the Global Distance Test–High Accuracy score (GDT-HA). Compared with oligo-GDTTS, oligo-GDTHA uses more stringent distance thresholds and is therefore more sensitive to fine-grained structural deviations[6].

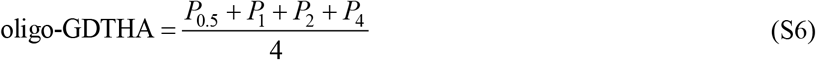

where *P*_0.5_, *P*_1_, *P*_2_ and *P*_4_ denote the fractions of aligned residue pairs whose corresponding Cα atoms are separated by no more than 0.5, 1, 2, and 4 Å, respectively, after structural superposition. A higher oligo-GDTHA value indicates greater overall structural accuracy of the candidate model across the entire complex.

##### DockQ-ave

DockQ provides an integrated assessment of the structural quality of protein-protein docking interfaces. It combines *F*_nat_, interface root-mean-square deviation (iRMSD), and ligand root-mean-square deviation (LRMSD), and is defined as[7]:

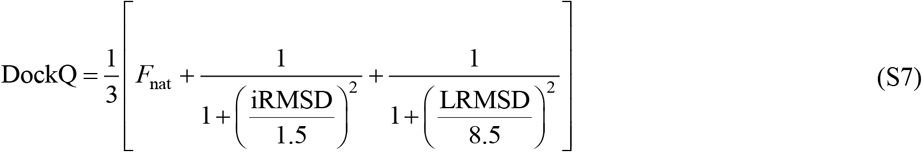

where *F*_nat_ is the fraction of native interfacial contacts correctly recovered in the candidate model; iRMSD is the root-mean-re deviation of interface residues after structural superposition; and LRMSD is the root-mean-square deviation of the d chain after superposition of the receptor chain. The constants 1.5 Å and 8.5 Å are the scaling factors used for iRMSD LRMSD, respectively, in the inverse-square scaling scheme of DockQ.

For complexes containing multiple interchain interfaces, DockQ-ave is the arithmetic mean of the DockQ scores over all matched interfaces and can be expressed as:

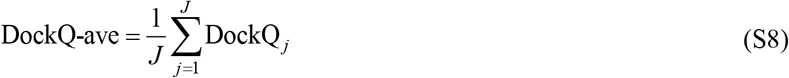

where *J* is the number of matched interfaces included in the evaluation, andDockQ *j* is the DockQ score for the *j* -th interface. A higher DockQ-ave value indicates better mean prediction quality across multiple inter-chain interfaces. In practice, chain mapping, the set of matched interfaces, and the aggregation procedure are determined by the evaluation program.

##### DockQ-wave

DockQ-wave is a weighted DockQ metric for protein complexes with multiple interfaces. Unlike DockQ ave, which assigns equal weight to all interfaces, DockQ-wave weights each interface-specific DockQ score by the number of native contacts in the corresponding interface of the reference structure. Its general form is[8]:

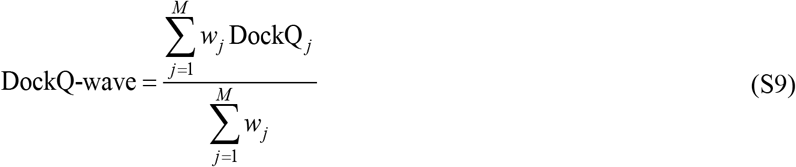

where *M* is the number of matched interfaces included in the evaluation, DockQ_j_ is the DockQ score for the *j* -th interface, and *j w* is the number of native inter-chain contacts in the corresponding interface of the reference structure. Thus, interfaces with more native contacts receive greater weight in the overall DockQ-wave score. A higher DockQ-wave value indicates better overall interface quality after accounting for the relative contributions of different interfaces.

##### IPS

IPS (Interface Patch Similarity) evaluates the agreement between the interface-residue patches of the candidate model he reference structure. It measures whether the candidate model correctly identifies the residues involved in interchain actions, without considering the specific residue–residue contact pairs. IPS is defined as the Jaccard similarity icient between the interface-residue sets of the two structures[9]:

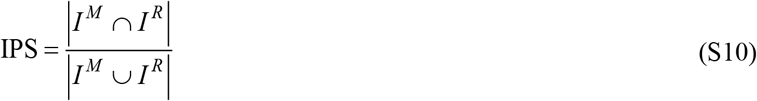

where *I*^*M*^ and *I*^*R*^ denote the sets of interface residues in the candidate model and the reference structure, respectively. An interface residue is defined as a residue participating in at least one inter-chain contact. Two residues from different chains are considered to be in contact if at least one pair of their heavy atoms is within 5 Å. IPS ranges from 0 to 1, with a higher value indicating greater agreement between the predicted and reference interface patches.

##### ICS

ICS (Interface Contact Similarity) evaluates the agreement between the interchain residue–residue contact patterns of the candidate model and the reference structure. Unlike IPS, which considers only whether individual residues belong to an interface, ICS retains the identities of the contacting residue pairs and therefore provides a more detailed assessment of interface-contact recovery.

Let *C*^*M*^ and *C*^*R*^ denote the sets of inter-chain residue contacts in the candidate model and the reference structure, respectively. Contact precision and contact recall are defined as:

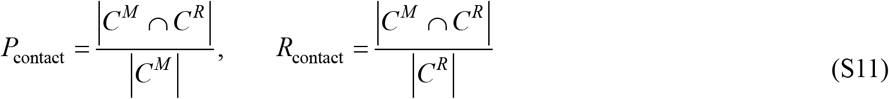

ICS is calculated as the F1-score of contact precision and contact recall[9]:

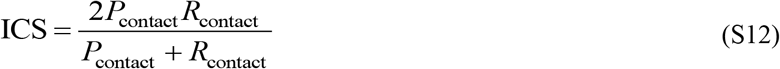

where an inter-chain contact is defined as a pair of residues from different chains for which at least one pair of heavy atoms is within 5 Å. ICS ranges from 0 to 1. A higher value indicates that the candidate model more accurately reproduces the native inter-chain contact pattern of the reference structure.

##### RMSD

RMSD (Root-mean-square deviation) measures the overall structural deviation between a predicted model and its reference structure based on the spatial distances between corresponding atoms after optimal rigid-body superposition. It is defined as:

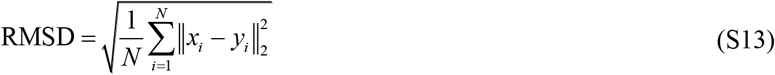

where *N* denotes the number of aligned atom pairs, and *i x* and *i y* denote the three-dimensional coordinates of the *i* -th corresponding atoms in the predicted model and the reference structure, respectively, after optimal superposition. RMSD is non-negative, with lower values indicating greater structural similarity between the predicted and reference structures.

#### Supplementary Note 2

Weighted penalty score

The weighted penalty score was calculated following the released implementation of the CASP16 QMODE3 evaluation framework[10]. The only difference between the Top-5 and Top-1 settings lies in the definition of the ranking error (RE). For the Top-5 setting, the ranking error for the OST metric *q* on target *t* was calculated as[10]:

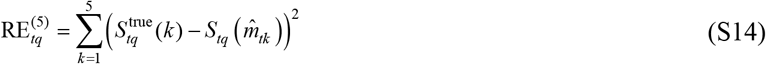

Where true 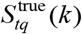 is the true OST score of the *k* -th model in the metric-specific reference Top-5 ranking, and 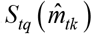 is the true OST score of the model selected by the method at the same position.

For the Top-1 setting, only one model was selected for each target. Instead of comparing against the reference Top-5 list, the ranking error was defined as the squared deviation between the selected model and the optimal score for that metric[10]:

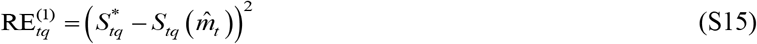

where 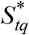 denotes the optimal score among all candidate models for target *t* under metric *q*.

For both settings, the metric-specific RE values were assembled into a RE vector[10],

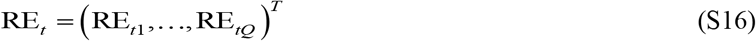

Where *Q* is the number of OST metrics included in the weighted-penalty calculation (*Q* = 9) for the oligomeric targets evaluated here, excluding RMSD, interface contact similarity (ICS), and interface patch similarity (IPS) from the 12 evaluated OST metrics).

The weighted penalty score was computed using the same covariance-weighted Mahalanobis distance defined in CASP16 QMODE3[10],

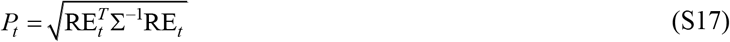

where ∑ denotes the covariance matrix indexed by the different OST metrics and ∑^−1^ denotes its Moore–Penrose pseudoinverse. The covariance matrix was calculated from OST scores of all evaluated models within each target category and was used to account for correlations among different metrics when combining ranking errors.

The resulting penalty scores were normalized within each target to account for differences in penalty score distributions among targets. Specifically, for each target, the penalty scores of all evaluated prediction groups were rescaled using min– max normalization[10]:

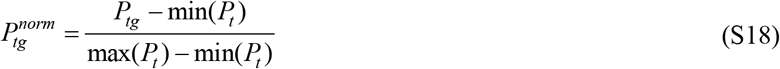

where *P*_*tg*_ denotes the weighted penalty score calculated for prediction group *g* on target *t*, and 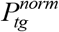 represents the corresponding normalized penalty score. The normalization was performed separately for each target, where the minimum and maximum values were calculated across all evaluated prediction groups within the same target. This procedure rescales the penalty scores for each target to a range between 0 and 1, allowing prediction groups evaluated on targets with different difficulty levels to be compared consistently. The final weighted penalty score was obtained by averaging the normalized penalty scores across all evaluated targets.

## Notes

### Competing Interest Statement

The authors have declared no competing interest.

